# Maternal defense against intruders changes her subsequent maternal behavior and neural circuitry

**DOI:** 10.64898/2026.06.30.735671

**Authors:** Patrese A. Robinson, Sandra Luz, Deep Patel, Gordon Barr, Seema Bhatnagar

## Abstract

Although female rats are typically less aggressive than male rats, lactating females will vigorously defend their nests and pups against an intruder. Much attention has been directed at the consequences of this aggression on the intruder and less on the consequences for the mother and her subsequent interactions with her pups. Here, we exposed resident Sprague-Dawley dams to the resident-intruder paradigm twice daily for five consecutive days, beginning when the dam’s (RES) pups were 7 days old, to assess social stress effects on maternal behavior and neurobiology. Controls were dams that had time-matched (TMC) separation from their pups but were not exposed to intruders, and naïve moms which were never separated nor exposed to an intruder (CTL). We assessed the dam’s subsequent behavior and interactions with her pups on Day 1 and Day 5, and Fos expression after Day 5 in select regions of the prefrontal cortex, amygdala, hypothalamus and periaqueductal gray of the midbrain. In separate cohorts, after pups were weaned, the dams underwent restraint stress and plasma corticosterone assayed. PCA analysis of the dam’s behaviors identified three components: normal self-focused behaviors; nurturing behaviors and rough non-nurturing behaviors. Relative to CTL, RES dams exhibited more disrupted behaviors towards their pups, including, rough transport, stepping on pups, and flinging/tossing pups around the cage. In contrast, TMC Dams showed some, but fewer changes relative to CTL, suggesting that separation from pups alone does not account for all disrupted behavior in RES dams. The bulk of these behavioral effects occurred in the first 5-10 min after reunion with the pups and were seen on both the first and fifth day of testing. Of the brain regions examined, the prefrontal cortex was activated by both the defeat/intruder stress (RES) and separation stress (TMC), whereas the dorsal PAG was activated specifically by the defeat/intruder stress. The medial and basolateral amygdala exhibited differential neuronal activity between the RES defeat/intruder-exposed dams and the other two groups. The RES moms exhibited an insufficient adrenocortical response to acute restraint stress. The results suggest that amygdala-dPAG activity is important for dissociating disrupted maternal care in RES (due to defense of the nest against an intruder) from simple pup separation, both of which activate the mPFC. The experience of repeatedly defending the nest may induce subsequent disruptions in HPA responses. The amygdala-dPAG pathway may regulate aspects of stress and emotional regulation exhibited by mothers who defend their offspring against intruders.

## Introduction

Stress-related mental illnesses, such as anxiety disorders, mood disorders, substance use disorders or eating disorders, can be exacerbated by myriad factors that impact the mother’s mental health during pregnancy, alter how the mother cares for her infants, reduce the quality of the mother-infant relationship, and increase the risk for poorer subsequent mental health outcomes for both infants and mothers ^1–3^. The importance of social influences on maternal stress levels has been recognized for decades, with early ^4,5^ and more recent ^6–10^ studies demonstrating an inverse relationship between the amount of maternal social support and reported levels of postpartum depression, stress, and anxiety.

Whereas community and social support can relieve stress in the mother and support her mental health, negative social experiences and peripartum social conflict can have outsized effects on maternal stress levels and wellbeing. During the prenatal period, social conflict is a strong predictor of maternal depression, even when levels of social support are high, suggesting that increased positive social experiences do not fully buffer against the effects of negative social experiences during this time ^10^. Relative to neutral experiences, negative social experiences for mothers increase physiological covariation (i.e., increased sympathetic activation in the dyad indicative of stress) between mothers and infants, more so than do positive experiences, even without the infant being present during the stressor ^11^. These acute negative social experiences in the mothers also produce infant stress-related behaviors such as social avoidance; in contrast, better quality maternal behavior is associated with less infant crying following reunion after maternal separation, regardless of the social experience valence ^11,12^. Importantly, these studies indicate that even for mothers with no documented mental health issues, acute negative social experiences can cause negative emotional responding in the infants. These reports demonstrate the inextricable connection between the maternal social experience, her peripartum mental health, and the mental health outcomes of her children.

To improve health outcomes for mothers and their infants experiencing peripartum/perinatal social stress, clinically relevant models that assess the neurobehavioral changes associated with social stress are needed. We developed a rodent model that induces social conflict by repeatedly exposing resident (RES) dams to female intruders which are defeated by the RES moms. In similar paradigms, during the postpartum period, social conflict increases stress-related behaviors in dams, such as excessive nesting, decreased nursing, and reduced pup grooming ^13–15^. The repeated defeat of intruders during social defeat stress shifts the profile of maternal behaviors: dams forgo direct maternal *care* during chronic social stress and engage in increased maternal *aggression,* vigilance, and time away from the nest, even when the social conflict occurs after the pups are removed and not in imminent danger ^13^. Although these reports reveal an effect of peripartum stress on mother behavior, they do not address the neurobehavioral mechanisms by which these social stressors alter maternal behavior or contribute to long-term mental health outcomes in the dams.

To accurately model the effects and outcomes of peripartum social stress in women, it is important to characterize neurobehavioral changes in preclinical models and to understand how underlying changes in maternal circuitry are reflected in their behavioral changes. The most complete models of maternal behavior recognize that the quality of rodent maternal behavior is determined by the balance of pup approach and withdrawal behaviors at any given point in the pup-dam interaction ^16–18^. Maternal behaviors towards pups naturally shift from approach during the early postnatal phase to withdrawal/avoidance, leading to weaning ^19^; maternal aggression also shifts as the maternal neuroendocrine environment changes during this window ^20–22^. Whereas the aforementioned rodent studies have suggested that dysregulation between approach, avoidance, and aggression can develop following maternal stress exposure, little has been done to fully characterize stress-related changes in maternal behavior and mental and/or emotional function or to model how these stressors alter the maternal neurobehavioral systems that control dam behaviors.

The circuit that typically coordinates maternal approach, avoidance, and aggression is extensive, with each neurobehavioral circuit sharing subregions or sending projections to the others. For maternal aggression, several hypothalamic nuclei are critical, as the hypothalamus receives information from the amygdala ^23–25^ that is integrated with other internal and external priming stimuli to facilitate appropriate aggressive responding ^26,27^. Further, in rodents, upon threat to the litter (typically an unfamiliar male), the vomeronasal organ (VMO) and main olfactory system send information specifically to the medial amygdala to integrate signals enabling maternal aggression ^28–30^. However, the amygdala also has overlapping contributions to maternal behaviors that support approach and withdrawal, which is coordinated through interactions of the mPFC with the medial amygdala (in the case of avoidance) and the BMA/BLA (during approach)^19^.

Early studies suggested that the mPFC may be responsible for the facilitation of maternal behavior because mPFC lesions delayed or disorganized the maternal responding to pups ^31–33^. Moreover, more recent reports suggest that the specific subregions of the mPFC, namely the IL and PL, perform executive control task that *facilitate* maternal responding early (in the case of the IL) and *reduce* maternal responding later (in the case of the PL) ^34–37^. Through its projections with the MPOA, olfactory system, dopaminergic, serotonergic, dorsal/ventral medial hypothalamic, and other regions, in addition to the amygdala, the mPFC facilitates the biphasic neurobiological maternal drive that vacillates across changing pregnancy stages^28,29,35,37^ (for current theoretical circuits, see^38,19,39^). Depending on the parturition stage, strong neurological input from an “approach system” or an “avoidance system” facilitates the switch in maternal response to pup stimuli as the pups mature ^19^.

In contrast to the mPFC, the PAG has a demonstrated role in maternal behavior that may drive behavioral choice away from pup-directed maternal behavior and toward vigilance/aggression that protects pups from threats or predatory behavior ^40,41^. Along its dorso-ventral/antero-posterior axis ^42,42–45^ and within its molecular neurochemistry ^43,44,46–50^, the PAG can both inhibit maternal care and facilitate maternal aggression or predatory behavior. The modulation of the behavioral switch and neurophysiological response is evident when dams are presented with conflicting motivation (i.e., exposure to predator odor while with pups versus without the pups ^49^), which demonstrates PAG’s role in managing motivational drives specifically in the lactating dam. Along with the amygdala, mPFC, other regions, activation in this expansive neural circuitry must be carefully coordinated across overlapping subsystems to ensure that the *appropriate* behaviors are produced in response to a particular stimulus (i.e., the pups versus an intruder).

There are many rodent models of postnatal stress, including maternal separation or deprivation, scarcity/low resource, and chronic social stress ^51–53^. However, these models typically focus on health outcomes in pups, with little exploration of the neurobiological changes that may persist in dams. For what little has been examined regarding dams following peripartum implementation of ELS protocols ^54,24,55,56^, these reports clearly demonstrate significant detrimental effects of peripartum stress on maternal health outcomes. Fortunately, these results reveal an opportunity to address a gap in our approach to modeling maternal peripartum stress effects, thereby providing clinically relevant profiles of neurobehavioral dysfunction that can aid in improving mental health outcomes in mothers. The current report contributes to this limited literature by implementing a rodent model of repeated social defeat stress in resident dams (RES) early in the postpartum period when nurturing behaviors predominate. After introducing an intruder into the home cage of the resident dam twice daily for 5 days, during which the dam typically attacked and defeated the intruder, we thoroughly assessed the dam’s maternal and non-maternal behaviors following the first day and fifth day of defeat and also measured changes in a subcircuit associated with coordinating maternal behaviors. We compared this social-aggression stress to the stress of removing the pups from the dam for an equivalent period (or not at all). As an initial exploration of long-term mental health outcomes, we also measured hormonal responding to restraint stress in the dams about two weeks after the end of social defeat stress and pups were weaned. Our results indicate that resident dams exhibit a range of maladaptive maternal behaviors following acute intruder exposure, characterized by decreased nurturing behaviors and increased abusive behavior. Importantly, some of these behavioral changes persist after chronic exposure. Further, in the mPFC-MeA-PAG subcircuit, distinct activation profiles distinguished cortical response to the intruder stress from pup-separation stress relative to control groups, and HPA activity was blunted following restraint. Our models is, to our knowledge, a novel approach to peripartum stress as it focuses on characterizing maternal (rather than pup) stress neurobiology by removing the pups during the intruder exposure and using smaller-sized female (rather than male) intruders which is common on other paradigms^13,15,57^. These changes remove imminent danger from the pups and reduce pup safety-related contributions to maternal neurobiological change, allowing us to better characterize maternal changes during a narrow window in which maternal behavior is strongest and neurobiological impacts are greatest. Further, our use of multiple control groups, extensive appraisal and characterization of maternal behavior via principle component analysis is, to our knowledge, the most extensive performed to date and provides a psychobiological scaffold on which to build the neurobiological framework of stress system dysregulation during the peripartum period.

## Methods

### Animals

Experimental subjects were adult female Sprague-Dawley rats obtained from Charles Rivers Laboratories. Timed-pregnant females, used as resident dams, arrived at gestation day 18, and virgin female Sprague-Dawley rats (200-225g) were used as intruders. Rats were individually housed in cages (cage dimensions 19” x 7.5” x 14") on a 12-h light, 12-h dark cycle (lights on at 0700) in a climate-controlled room with ad libitum food and water. Rats were given 5 days of acclimation before experimentation. Studies were approved by Children’s Hospital of Philadelphia Research Institute’s Institutional Animal Care and Use Committee and conformed to the National Institutes of Health Guide for the Use of Laboratory Animals.

### Social defeat stress procedure

The social defeat procedure was conducted from postnatal day (PD)7-11. The social defeat paradigm used in this study was based on the resident-intruder model ^58–60^ starting at approximately 0800h to 1300h each day (see Figure 1). Sprague-Dawley rats were randomly assigned to one of three experimental conditions: Control [Naive-retrieval (CTL)], Time-Matched to resident Controls, (TMC) or social defeat Resident (RES) group. The duration of the treatment for all groups was two 30-minute consecutive sessions for five days. On Day 1 and Day 5 only, for the CTL group, immediately before recording maternal observations, pups were gently removed from the nest and scattered around the cage, but were never taken away from the dam. On Days 1-5, for RES prior to social defeat and for the TMC group, the litter was gently removed and placed in a bucket with bedding on a heating pad. TMC dams were then undisturbed. In the RES group, a smaller intruder was placed into the home cage of the resident dam for each of the 5 consecutive days, ending on PD11. Characteristically, the resident and intruder investigate each other for a short period (1–3 min), followed by an attack by the resident that ends in the defeat of the intruder. A defeat is pronounced when the intruder displays a supine posture and freezes for 2–3 s. The latency to defeat was then recorded. The resident and intruder are then separated by a wire mesh barrier until 30 min elapses from the time of initial placement of the intruder into the resident’s cage. The barrier allows for visual, auditory, and olfactory contact but prevents any further physical contact. As a way to minimize risk for excessive injury, wire mesh separation also occurs after 5 attacks. If no defeat occurs within 15 min (900 s), the rats are separated with a wire mesh barrier for the remaining 15 min; in this case, the defeat latency is entered as 900 s. This process was repeated one more time for 60 minutes total of social defeat exposure. A new intruder was used for the second social defeat exposure. Intruders were rotated so that they were not used more than twice per day and no resident defeated the same intruder more than once times over the 5 days. CTL and TMC rats were not exposed to this defeat paradigm but had a divider added to their home cage for the same two 30-minute windows, while RES rats underwent the defeat procedures.

**Figure 1:**
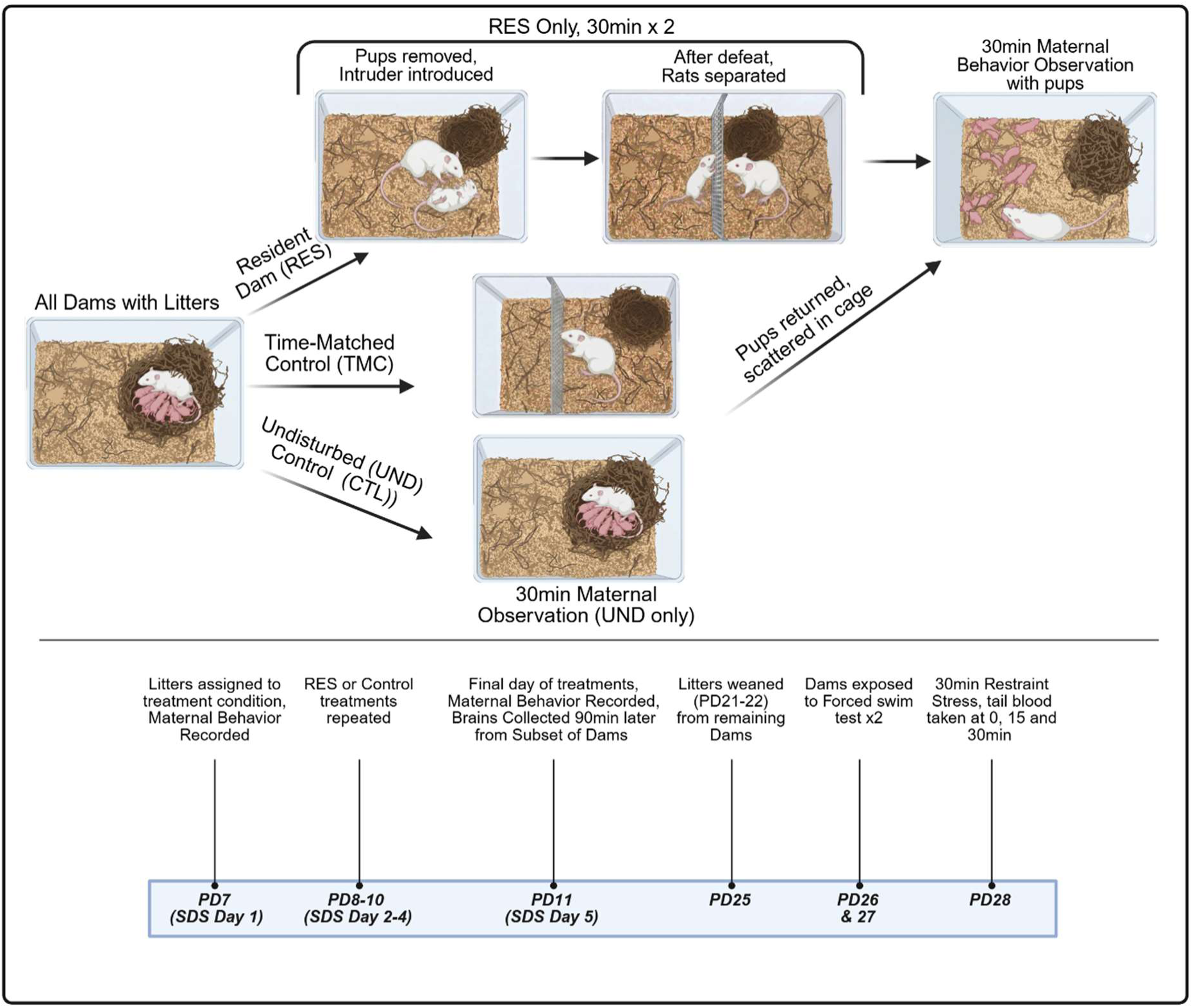
This figure illustrates the social stress manipulation and timeline of experimental procedures used in the current report. One set of dams was euthanized at the end of SDS exposure, and a different subset were retained and subjected to restraint stress.

### Maternal Observations

Observations of dams and their nests were video recorded using an overhead camera through plexiglass and Ethovision XT (version 13, Noldus) software. Following removal of the intruder from the RES cage, the divider was removed and, for the CTL, TMC, and RES groups, pups were gently scattered outside of the nest immediately prior to the start of recording. To collect reference recordings for baseline maternal behavior, a subset of CTL (n=4, prior to being scattered) and a set of undisturbed dams (n=4) were observed [Naïve-Undisturbed (UND)] and recorded for 30min with no disturbance; these data are displayed in Figures 2-7 and Supplemental Figures 2-3 for illustrative purposes only and were not statistically compared to the three experiment groups. Observations lasted 30min and behaviors were assessed for their total duration (s) and frequency (#) following Day 1 (Test 1) and Day 5 (Test 2) of the social defeat paradigm (see Table 1 for behavioral descriptions).

**Figure 2:**
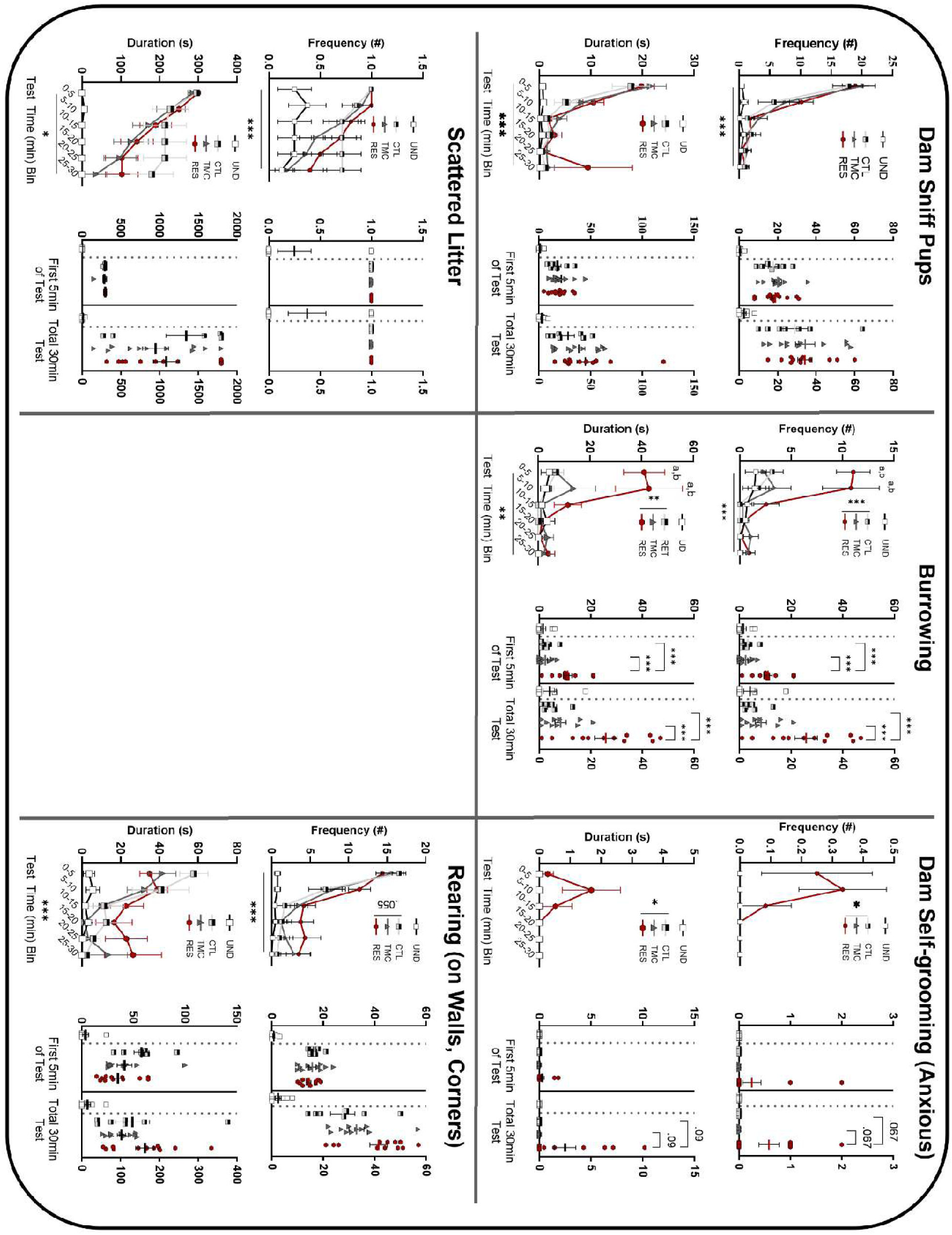
Mean (±SEM) Duration (s) and Frequency (#) of maternal behaviors from RC1 on Day 1 (Test 1) of social defeat stress or control conditions. 2-way repeated measure ANOVA for (Time bin x Condition), significant interaction with Holm-Šídák test post hoc pairwise comparisons represented as: ^a^RES ≠TMC, ^b^RES ≠CTL, and ^c^TMC ≠CTL, *p < .05, **p < .01, or ***p < .001

**Figure 3:**
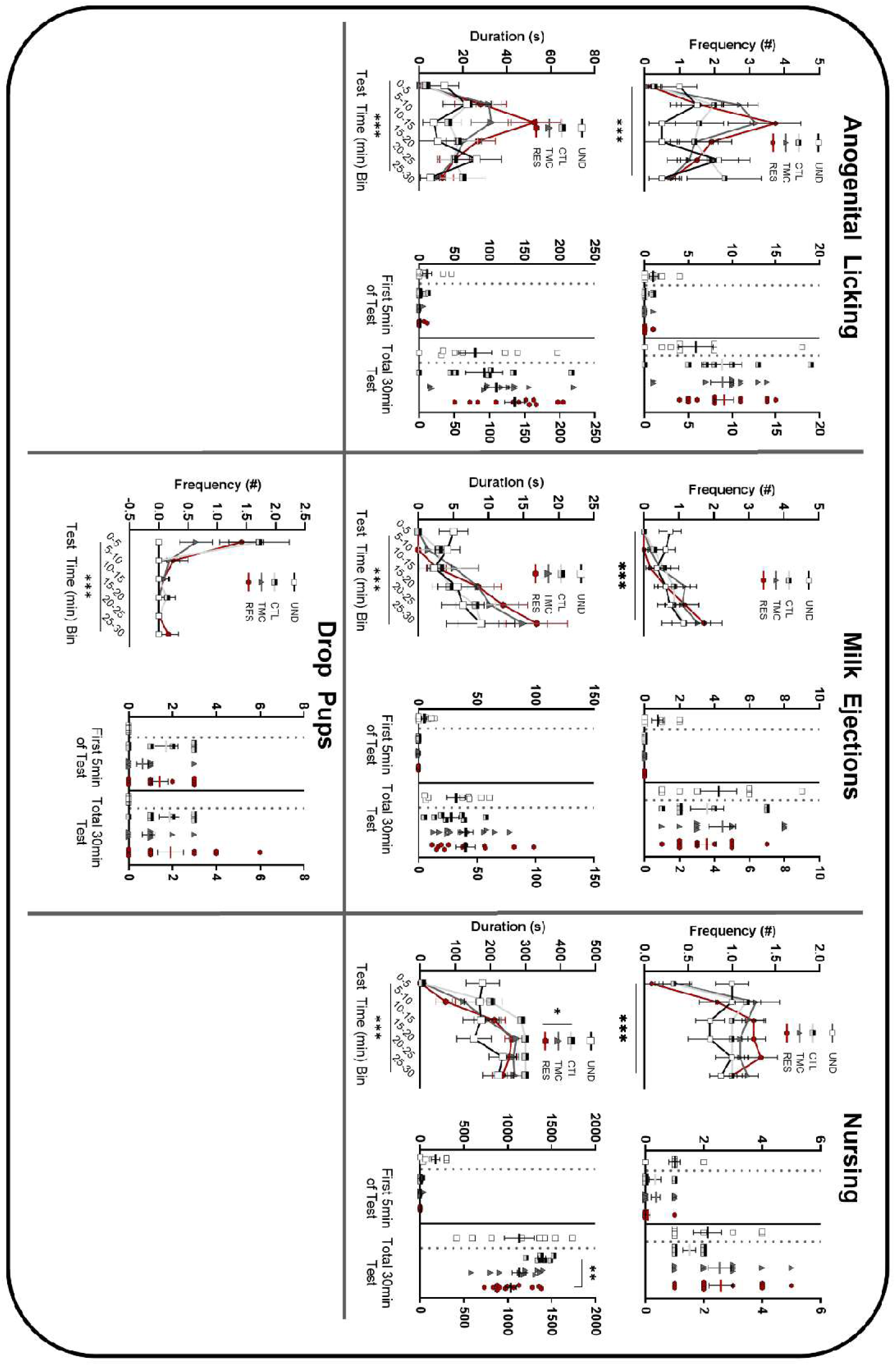
Mean (±SEM) Duration (s) and Frequency (#) of maternal behaviors from RC2 on Day 1 (Test 1) of social defeat stress or control conditions. 2-way repeated measure ANOVA for (Time bin x Condition), significant interaction with Holm-Šídák test post hoc pairwise comparisons represented as: ^a^RES ≠TMC, ^b^RES ≠CTL, and ^c^TMC ≠CTL, *p < .05, **p < .01, or ***p < .001.

**Figure 4:**
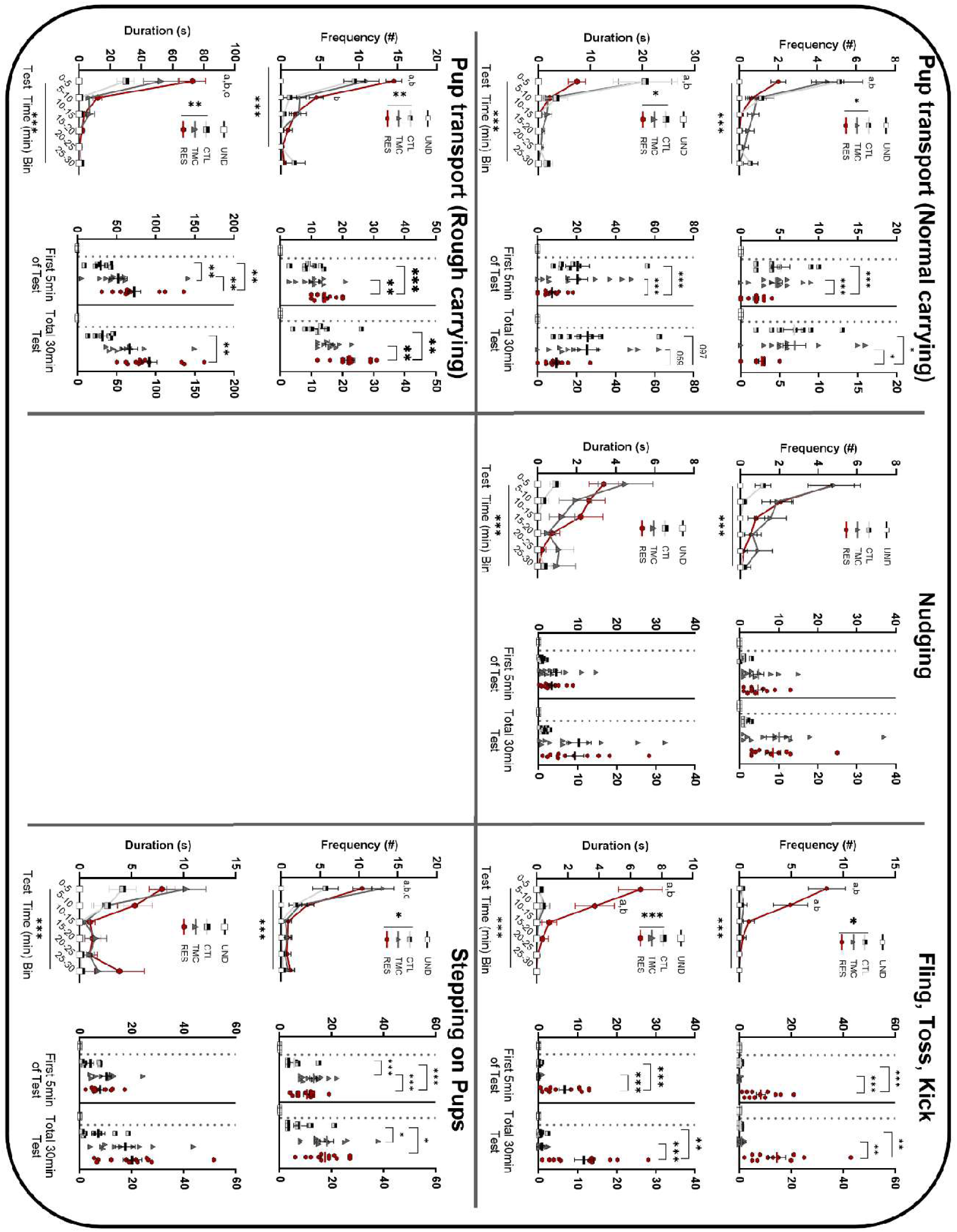
Mean (±SEM) Duration (s) and Frequency (#) of maternal behaviors from RC3 on Day 1 (Test 1) of social defeat stress or control conditions. 2-way repeated measure ANOVA for (Time bin x Condition), significant interaction with Holm-Šídák test post hoc pairwise comparisons represented as: ^a^RES ≠TMC, ^b^RES ≠CTL, and ^c^TMC ≠CTL, *p < .05, **p < .01, or ***p < .001

**Figure 5:**
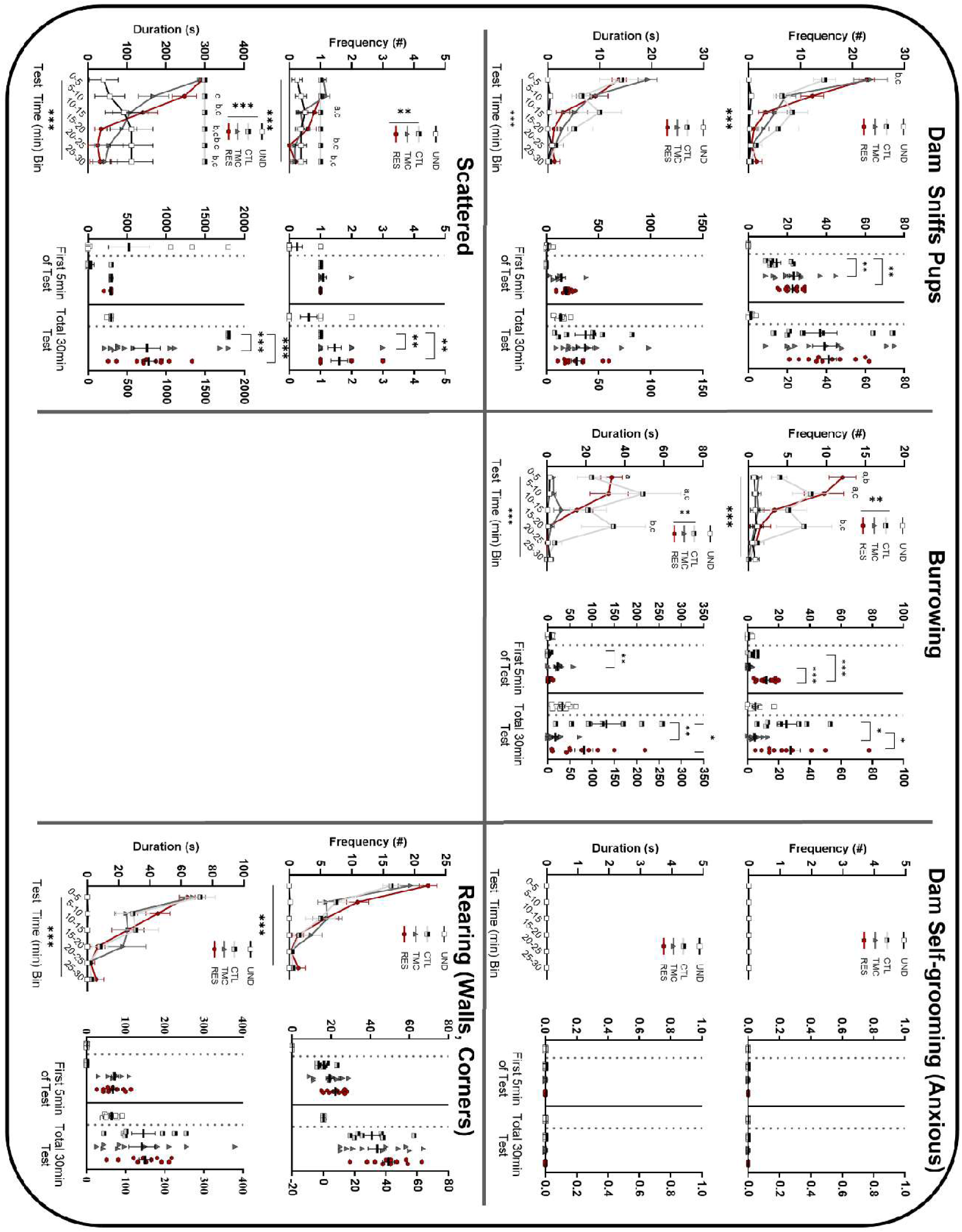
Mean (±SEM) Duration (s) and Frequency (#) of maternal behaviors from RC1 on Day 5 (Test 2) of social defeat stress or control conditions. 2-way repeated measure ANOVA for (Time bin x Condition), significant interaction with Holm-Šídák test post hoc pairwise comparisons represented as: ^a^RES ≠TMC, ^b^RES ≠CTL, and ^c^TMC ≠CTL, *p < .05, **p < .01, or ***p < .001

**Figure 6:**
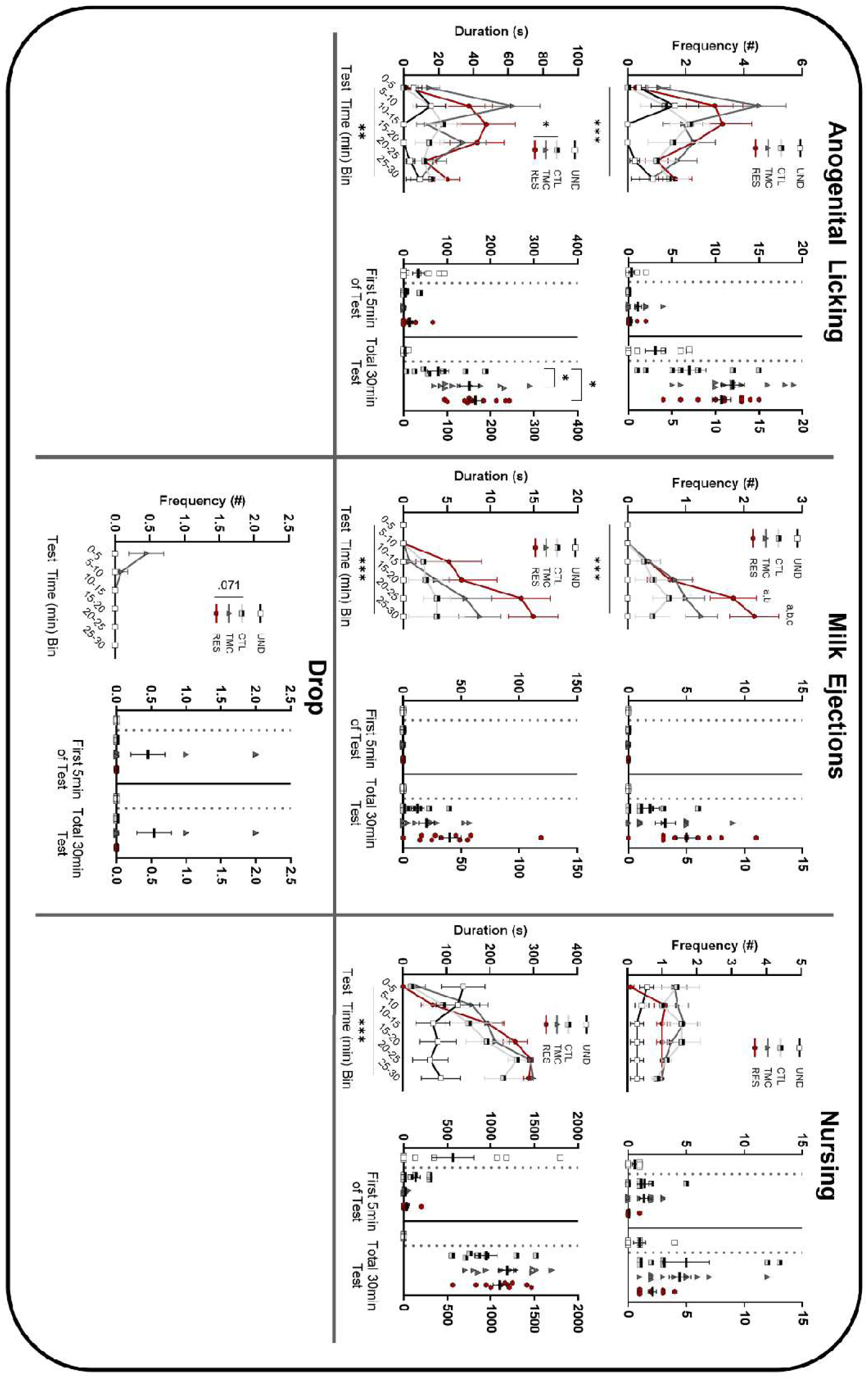
Mean (±SEM) Duration (s) and Frequency (#) of maternal behaviors from RC2 on Day 5 (Test 2) of social defeat stress or control conditions. 2-way repeated measure ANOVA for (Time bin x Condition), significant interaction with Holm-Šídák test post hoc pairwise comparisons represented as: ^a^RES ≠TMC, ^b^RES ≠CTL, and ^c^TMC ≠CTL, *p < .05, **p < .01, or ***p < .001.

**Figure 7:**
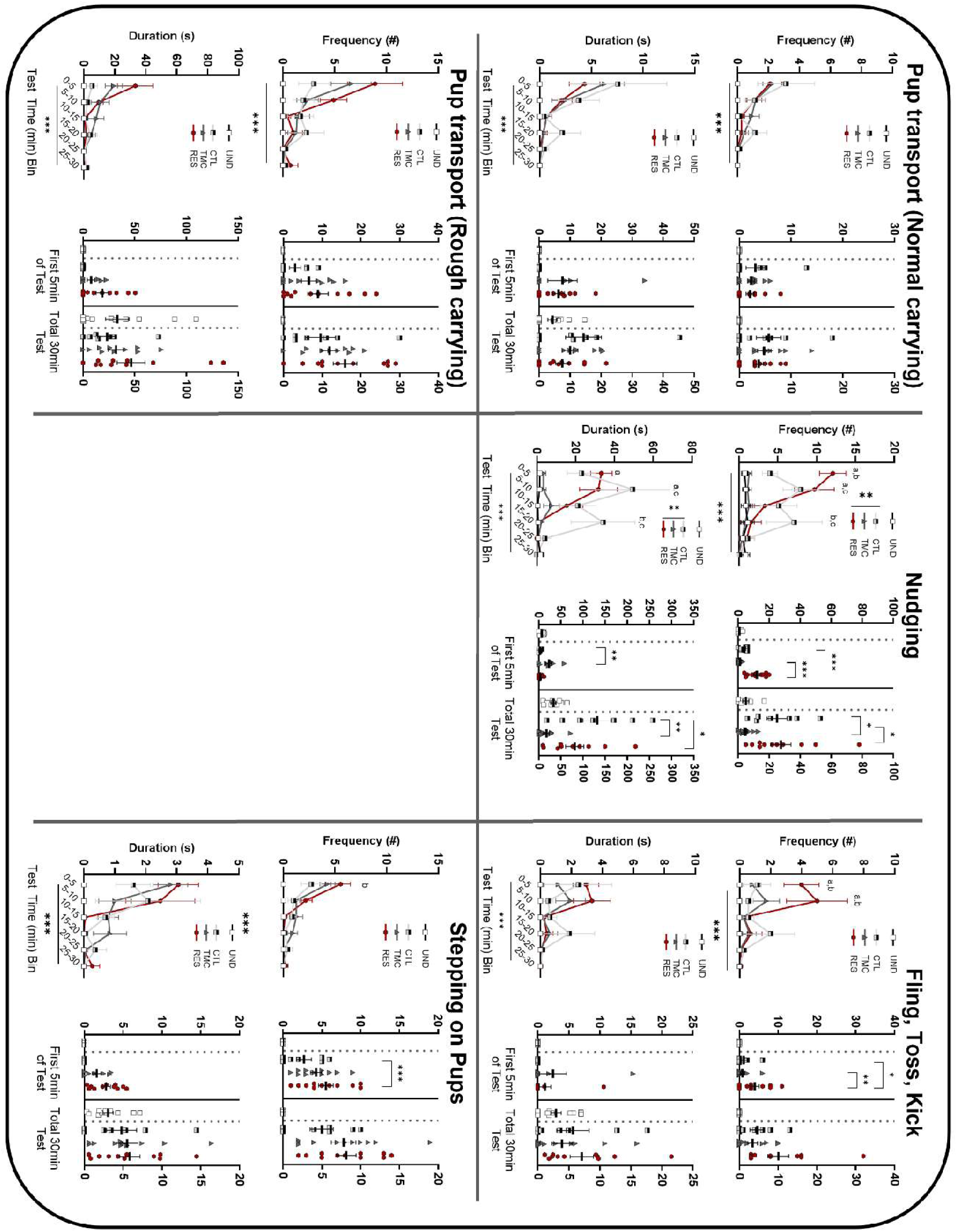
Mean (±SEM) Duration (s) and Frequency (#) of maternal behaviors from RC3 on Day 5 (Test 2) of social defeat stress or control conditions. 2-way repeated measure ANOVA for (Time bin x Condition), significant interaction with Holm-Šídák test post hoc pairwise comparisons represented as: ^a^RES ≠TMC, ^b^RES ≠CTL, and ^c^TMC ≠CTL, *p < .05, **p < .01, or ***p < .001

**Table 1.**
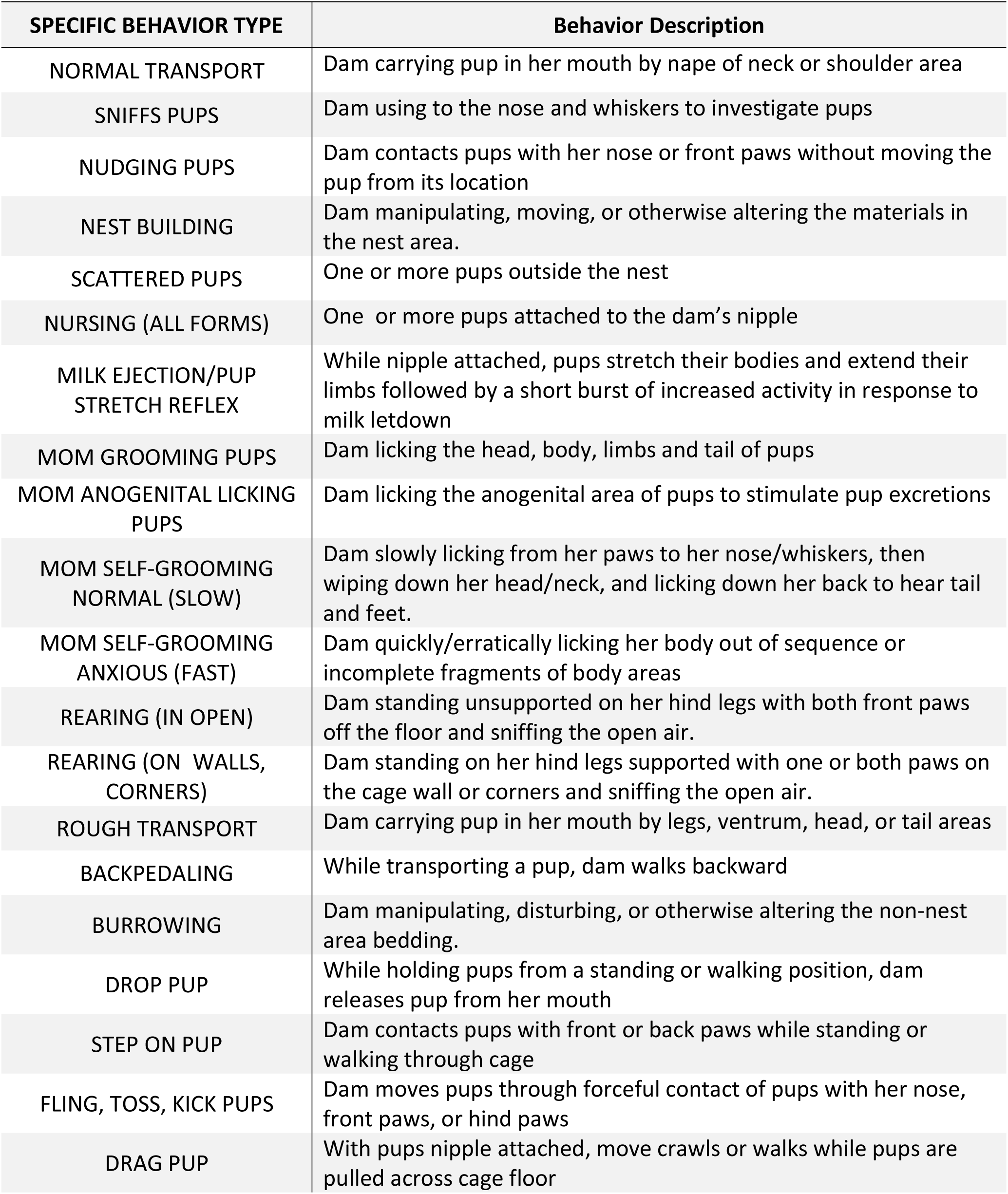
Description of Maternal Behaviors Measured following the Social Defeat Paradigm.

| **SPECIFIC BEHAVIOR TYPE** | **Behavior Description** |
| --- | --- |
| NORMAL TRANSPORT | Dam carrying pup in her mouth by nape of neck or shoulder area |
| SNIFFS PUPS | Dam using to the nose and whiskers to investigate pups |
| NUDGING PUPS | Dam contacts pups with her nose or front paws without moving the pup from its location |
| NEST BUILDING | Dam manipulating, moving, or otherwise altering the materials in the nest area. |
| SCATTERED PUPS | One or more pups outside the nest |
| NURSING (ALL FORMS) | One or more pups attached to the dam's nipple |
| MILK EJECTION/PUP STRETCH REFLEX | While nipple attached, pups stretch their bodies and extend their limbs followed by a short burst of increased activity in response to milk letdown |
| MOM GROOMING PUPS | Dam licking the head, body, limbs and tail of pups |
| MOM ANOGENITAL LICKING PUPS | Dam licking the anogenital area of pups to stimulate pup excretions |
| MOM SELF-GROOMING NORMAL (SLOW) | Dam slowly licking from her paws to her nose/whiskers, then wiping down her head/neck, and licking down her back to hear tail and feet. |
| MOM SELF-GROOMING ANXIOUS (FAST) | Dam quickly/erratically licking her body out of sequence or incomplete fragments of body areas |
| REARING (IN OPEN) | Dam standing unsupported on her hind legs with both front paws off the floor and sniffing the open air. |
| REARING (ON WALLS, CORNERS) | Dam standing on her hind legs supported with one or both paws on the cage wall or corners and sniffing the open air. |
| ROUGH TRANSPORT | Dam carrying pup in her mouth by legs, ventrum, head, or tail areas |
| BACKPEDALING | While transporting a pup, dam walks backward |
| BURROWING | Dam manipulating, disturbing, or otherwise altering the non-nest area bedding. |
| DROP PUP | While holding pups from a standing or walking position, dam releases pup from her mouth |
| STEP ON PUP | Dam contacts pups with front or back paws while standing or walking through cage |
| FLING, TOSS, KICK PUPS | Dam moves pups through forceful contact of pups with her nose, front paws, or hind paws |
| DRAG PUP | With pups nipple attached, move crawls or walks while pups are pulled across cage floor |

### Defeat-induced neuronal activity in select brain regions

In a sub-cohort of experimental animals, brains were collected at 60 min after the end of the last defeat exposure on day 5. Brains were frozen until sectioned at 20 µm and immunostained for cFos (1:1250, sc-52; Santa Cruz Biotechnology, Santa Cruz, CA). Neuronal activation in select brain regions (see Figures 8-9, Supplemental Figures 4-5) implicated in stress responses and/or defensive responses was assessed and analyzed in NIH Image J by two investigators blind to treatment conditions.

**Figure 8:**
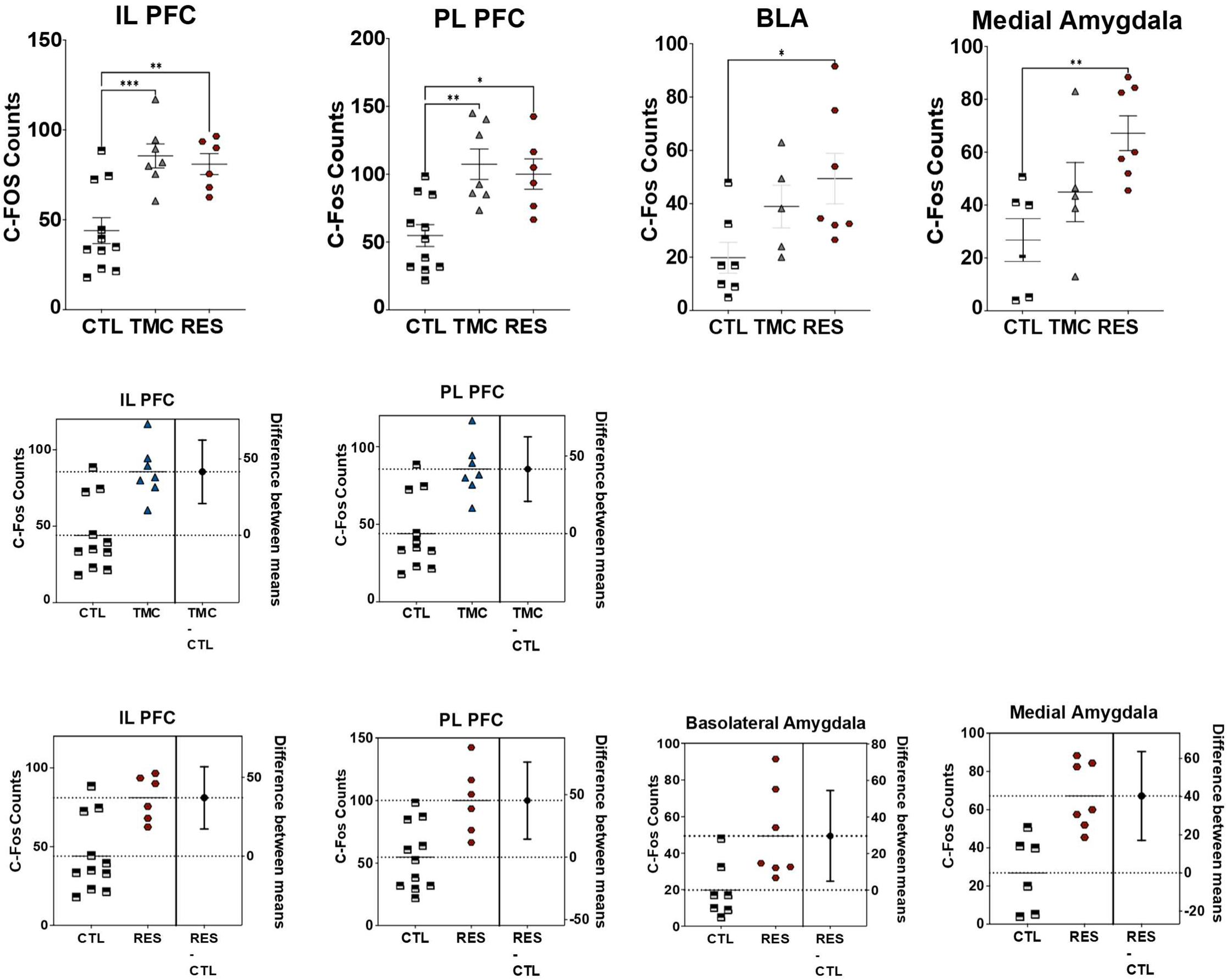
Mean (±SEM) count of c-Fos expression cells in select regions of the brain. (Top row) The group sizes for the CTL, TMC, and RES dam groups for each brain region is as follows: for PL and IL mPFC n=11, 7, and 6; MEA n=6, 5, and 7. *indicates p < .05, **p < .01, ***p < .001. The estimation plots (rows 2 and 3) presents the data points (left panel) and mean difference between the two conditions plus/minus the standard deviation (right panel) for a single behavior.

**Figure 9:**
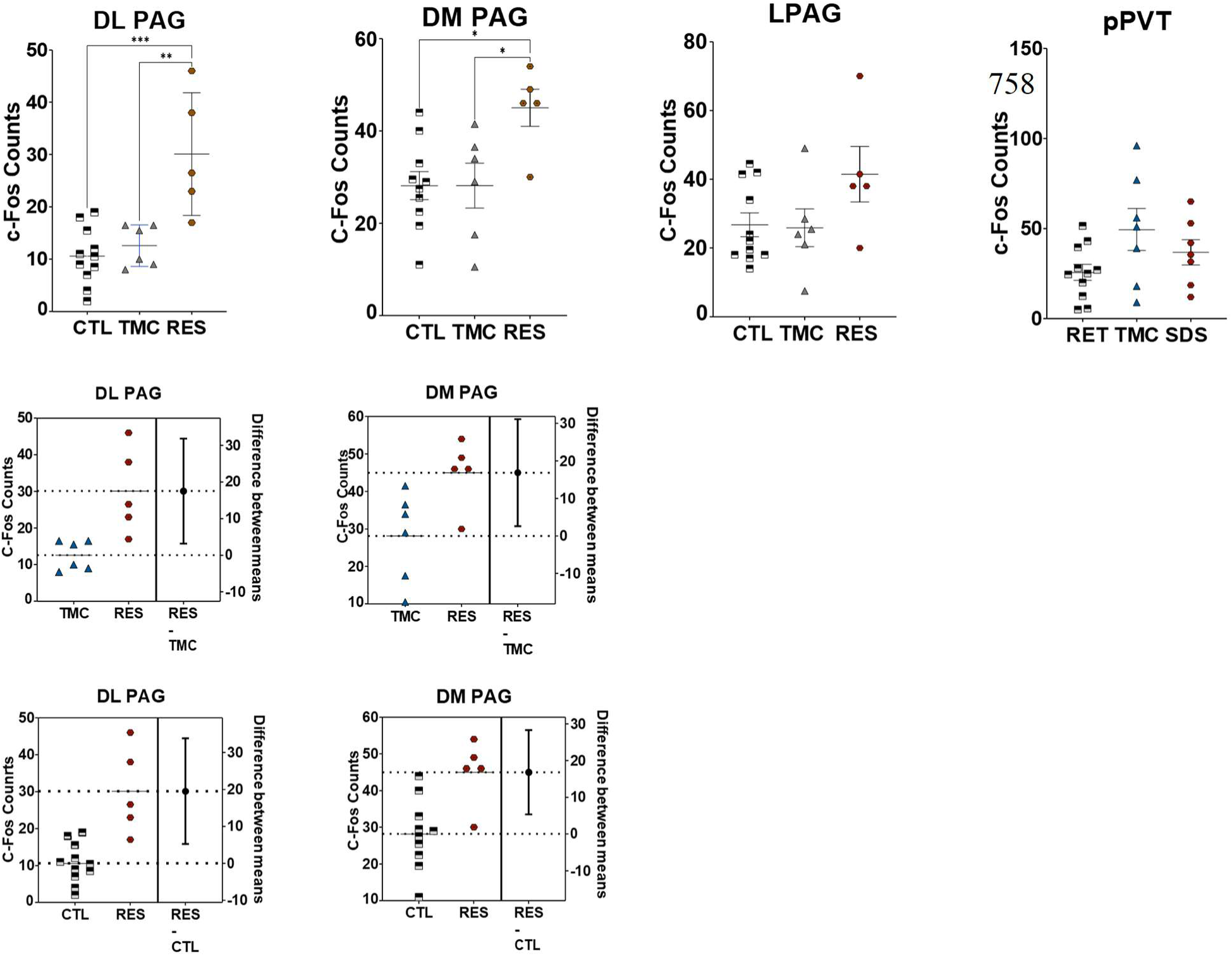
Mean (±SEM) count of c-Fos expression cells in select regions of the brain. (Top row) The group sizes for the CTL, TMC, and RES dam groups for each brain region is as follows: PVT n=11, 7, and 7 and for all PAG subregions n=11, 6, and 5 plots (rows 2 and 3) presents the data points (left panel) and mean difference between the two conditions plus/minus the standard deviation (right panel) for a single behavior.

### Forced swim test and Corticosterone responses to restraint

A separate group of dams was retained through pup weaning on PD25 and administered the forced swim test on PD26 and 27. Rats were placed in a glass cylinder filled with 60 cm of water so that their tails could not touch the bottom of the cylinder while floating. Rats underwent a 15 min training phase (PD26) followed by a 5 min test phase on the following day (PD27). The test phase was videotaped and coded for time (s) immobile, swimming, and climbing behaviors by two coders who were blind to the experimental conditions.

One day after the forced swim tests, on PD28, dams were exposed to 30 minutes of restraint to determine hypothalamic-pituitary-adrenal (HPA) activity in response to acute stress. Tail blood (via nicking of the tail vein) was taken at 0 minutes (as soon as the animal was removed from the home cage and placed in a plastic tubular restrainer), again at 15 minutes and 30 minutes (during restraint). At 60 min after the start of restraint, the animal was removed from its home cage and trunk blood collected (recovery time point) after rapid decapitation. Blood was centrifuged at 2500 rpm for 15 minutes. Plasma was aliquoted into new Eppendorf tubes and stored at -80°C. Plasma corticosterone was assayed with a Radioimmunoassay kit from MP Biomedical (Orangeburg, NY). The minimum levels of detection for corticosterone were 0.6μg/dl. Intra- and inter-assay variability was less than 10%. The area under the curve for hormone levels at these four time points was then calculated as the integrated hormone response to restraint.

## Statistics

### Principal Component Analysis of Maternal behaviors

To help organize the multiple behaviors into a more coherent structure, we performed a Principal Component Analysis of the combined frequency and duration of each behavior (ProMax oblique rotation; JASP [JASP Team 2026; (https://jasp-stats.org); Version 0.95.4]. We did not define *a priori* the number of components, which were determined by the results of the Scree analysis (Supplemental Figure 1) comparing PC loading from the actual data to the same data randomized and identified three components (Table 2). Different analysis parameters (e.g. orthogonal rotation; different oblique rotations; unrotated solution) produced similar results.

**Table 2.** Component Loadings for Maternal Behaviors.

|  | **RC1** | **RC2** | **RC3** | **Uniqueness** |
| --- | --- | --- | --- | --- |
| BURROWING | 0.795 |  |  | 0.386 |
| DRAG PUP | 0.740 |  |  | 0.476 |
| SCATTERED LITTER | 0.711 |  |  | 0.322 |
| REARING (WALLS AND CORNERS) | 0.705 |  |  | 0.300 |
| MOM SELF-GROOMING NORMAL (SLOW) | 0.645 |  | -0.498 | 0.443 |
| MOM SELF-GROOMING ANXIOUS (FAST) | 0.627 |  |  | 0.599 |
| SNIFFS PUPS | 0.577 |  |  | 0.407 |
| REARING (OPEN) | 0.543 |  |  | 0.665 |
| NURSING (ALL FORMS) |  | 0.859 |  | 0.184 |
| MOM ANOGENITAL LICKING PUPS |  | 0.836 |  | 0.248 |
| MOM GROOMING PUPS |  | 0.788 |  | 0.371 |
| MILK EJECTION/PUP STRETCH REFLEX |  | 0.622 |  | 0.628 |
| DROP PUP |  | -0.531 |  | 0.540 |
| STEP ON PUP |  | -0.491 | 0.587 | 0.402 |
| NUDGING PUPS |  |  | 0.796 | 0.311 |
| ROUGH TRANSPORT |  |  | 0.734 | 0.328 |
| MOM BACKPEDALING |  |  | 0.719 | 0.471 |
| FLING, TOSS, KICK PUPS |  |  | 0.566 | 0.627 |
| NORMAL TRANSPORT |  |  | 0.566 | 0.533 |
| NEST BUILDING |  |  |  | 0.618 |

*Note. Applied oblique rotation method is Promax.*

### Maternal behavior observations

We analyzed video recordings for the following number of litters per group: UND n=8, CTL n=7, TMC n=11, RES n=12. Outlier performance per behavioral measure was determined using GraphPad Prism ROUT method of identification and the Q coefficient set to 1%, per program recommendations (between 0-3 outliers removed per group). The duration (s) and frequency (#) of behaviors performed by dams was scored offline by observers blind to their experimental condition. Two-way repeated measures ANOVA were used to assess the effects of maternal condition (condition: CTL, TMC, RES; between-subjects) on dam behavior across six time bins over the course of the 30min test (time: 0-5min, 5-10min, 10-15min, 15-20min, 20-25min, 25-30min; within-subjects). Statistics (see Tables 3-6) regarding non-significant effects and the main effects for time are omitted for brevity. UND behaviors were at floor or ceiling levels with no variability in most analyses, so their data are depicted in the associated figures for illustrative purposes only but were not included in the statistical analyses. Significant effects were followed up with the Holm-Šídák post-hoc test. Following the ANOVA and pairwise post hoc comparisons, we calculated estimation plots to provide more information about effect size ^61^ using Prism GraphPad (v11.02 MacOS, Figure 8-9, Supplemental Figures 4-5). The left panel of each estimation graph shows the individual data points and the mean of each. The right panel shows the difference between the two groups ± one SD.

**Table 3.** Summary of 2-way repeated measures ANOVA results from Day 1 (Test 1): Duration (s)

| Maternal Behavior:<br>Test 1 Duration (s) | 5min Time x<br>Condition<br>Significance |  |  |  | 30min Condition<br>Significance |  |  |  |
| --- | --- | --- | --- | --- | --- | --- | --- | --- |
|  |  | CTL v<br>TMC | CTL v<br>RES | TMC v<br>RES |  | CTL v<br>TMC | CTL v<br>RES | TMC v<br>RES |
| Burrowing | <.001 |  | ✓✓✓ | ✓✓✓ | 0.004 |  | ✓ | ✓ |
| Drag pup |  |  |  |  |  |  |  |  |
| Scattered Litter |  |  |  |  |  |  |  |  |
| Rearing (Walls and Corners) |  |  |  |  |  |  |  |  |
| Mom self-grooming anxious (fast) |  |  |  |  | 0.046 |  | 0.099 | 0.099 |
| Mom self-grooming normal (slow) |  |  |  |  |  |  |  |  |
| Sniffs pups |  |  |  |  |  |  |  |  |
| Rearing (open) |  |  |  |  |  |  |  |  |
| Nursing (all forms) |  |  |  |  | 0.011 | 0.053 | ✓✓ |  |
| Mom anogenital licking pups |  |  |  |  |  |  |  |  |
| Mom grooming pups |  |  |  |  |  |  |  |  |
| Milk Ejection/pup stretch reflex |  |  |  |  |  |  |  |  |
| Drop Pup (Negative) |  |  |  |  |  |  |  |  |
| Nudging pups |  |  |  |  |  |  |  |  |
| Rough transport | <.001 | ✓✓✓ | ✓✓✓ | ✓✓✓ | 0.002 | 0.078 | ✓✓ | 0.078 |
| Mom Backpedaling |  |  |  |  |  |  |  |  |
| Step on pup |  |  |  |  | 0.060 |  |  |  |
| Fling, Toss, kick pups | <.001 |  | ✓✓ | ✓✓ | <.001 |  | ✓✓ | ✓✓✓ |
| Normal transport | 0.002 |  | ✓✓✓ | ✓✓✓ | 0.038 |  | 0.078 | 0.069 |
| Nest Building |  |  |  |  |  |  |  |  |

Note: This table illustrates, for Test 2 Duration (s) of Maternal behaviors, the p-values for significant (highlighted in green) or trending ( $p < .10$ ) interaction (5min Time x Condition column) or main (30min Condition column) effects. The far-left column lists the measured maternal behaviors with components 1-3 shaded as described below. The middle column shows the p-values for the 5min Time x Condition interaction effect. The main effect of time is represented on the bar graphs for each behavior in Figures 2-7 and Supplementary Figures 2-3. The far-right column shows p-values for the condition main effect. Check marks indicate the level of significance as determined by Holm-Šidák test post hoc analysis of pairwise comparisons and described below.
RC 1 $p < .05$ ✓
RC 2 $p < .01$ ✓✓
RC 3 $p < .001$ ✓✓✓

### Corticosterone Assays

Plasma corticosterone (Figure 10) was analyzed by two-way ANOVAs (time: 0min, 15min, 30min, 60min, within-subjects x condition), integrated corticosterone was analyzed by one-way ANOVA.

**Figure 10:**
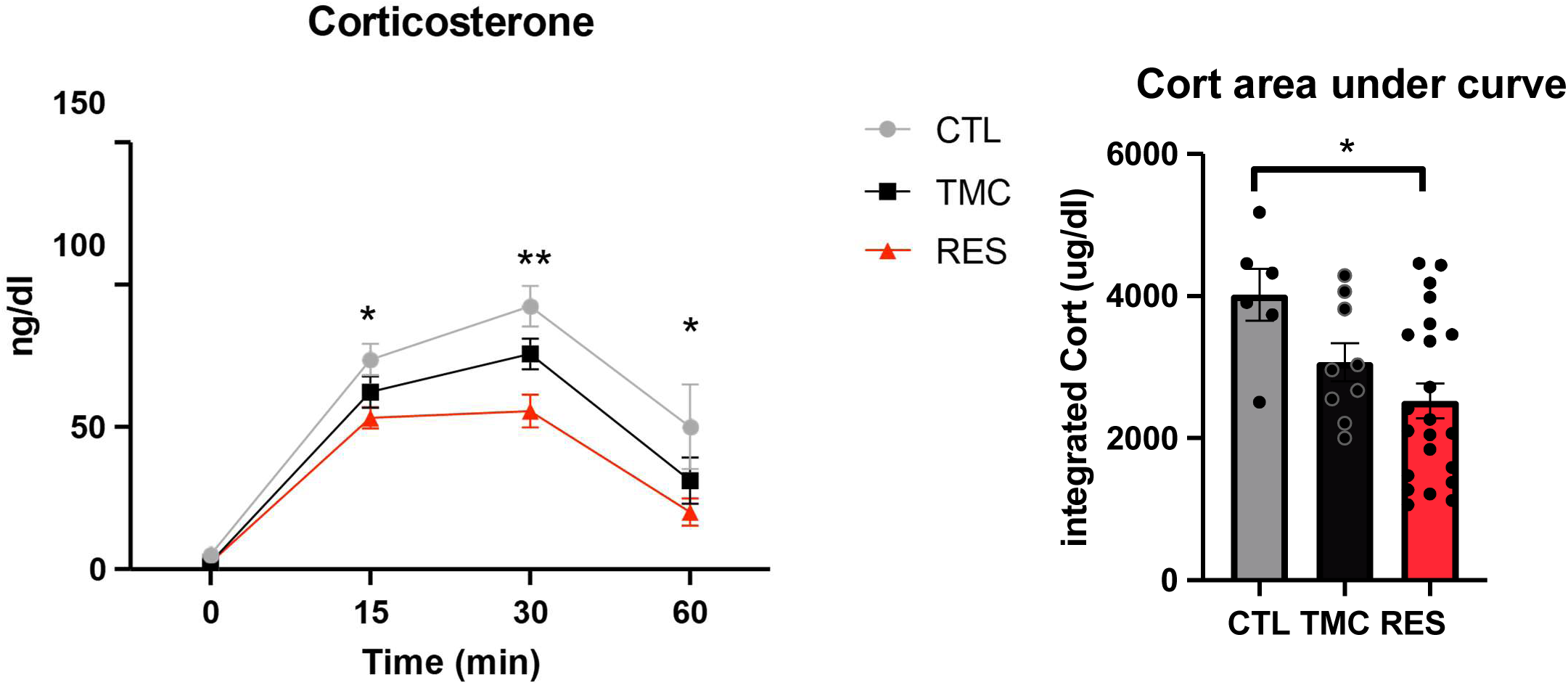
Corticosterone levels in dams following post-weaning Restraint Stress test. (Left panel) Mean (±SEM) amounts (ng/dL) CORT levels at different time points after restraint test. (Right panel) Integrated corticosterone analyses comparing overall levels of CORT expression between groups. Two-way ANOVA *indicates p < .05, **p < .01. One-way ANOVA bracketed “[“ *indicates p < .05, **p < .01.

## Results

### Principal Component Analysis for Maternal Observations

The PCA showed that three components had eigen values above those of the simulated parallel analysis (Supplemental Figure 1). We chose to focus on these three components which accounted for about 58% of the variance. The loadings of the individual components are in Table 2. Component 1 (RC1) roughly corresponded to self-focused and exploratory behaviors but with some pup interaction (sniffing pups, dragging pups). Component 2 (RC2) represented more positive mom-pup interactions related to maternal care with positive loadings such as nursing with milk ejection, anogenital licking, grooming pups, and negative loadings for stepping on the pups. Component 3 (RC3) contains largely disrupted interactions between the pups and dam. Positive loadings include, for example, rough transport, stepping on pup, tossing pups, scattering pups among others; negative loading was Mom slowly self-grooming. Subsequent data presentation is roughly organized around these groupings.

### Maternal Behavioral Outcomes

The main effects and interaction effects are summarized for both duration and frequency of behaviors on Day 1 (Tables 3 and 4, respectively) and Day 5 (Tables 5 and 6, respectively). Most of the effects that are significant occur in the RC3 related behaviors for both frequency and duration. More changes occur on the first day of testing than on the fifth day, suggesting some form of adaptation or habituation.

**Table 4.**
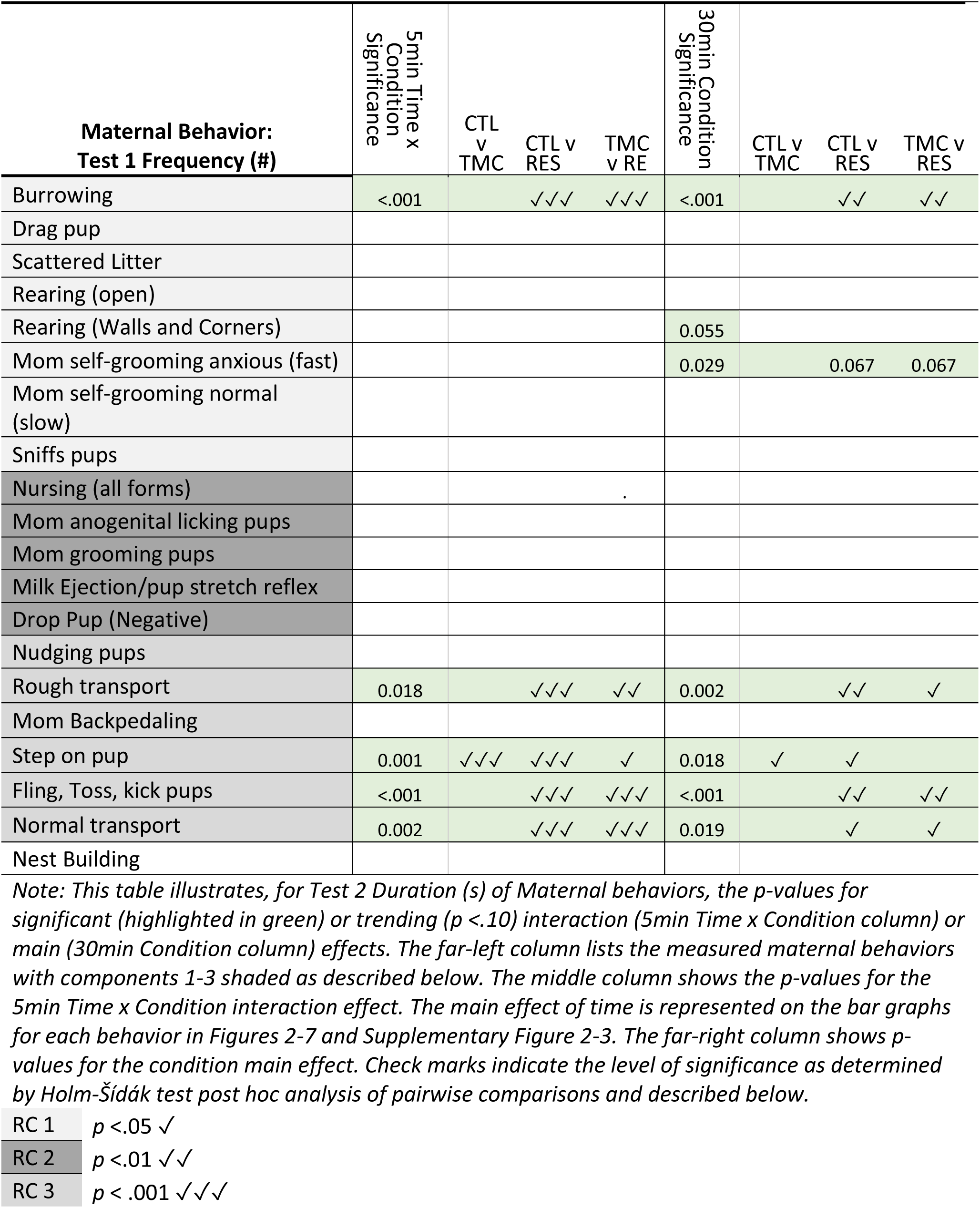
Summary of 2-way repeated measures ANOVA results from Day 1 (Test 1): Frequency (#)

**Table 5.**
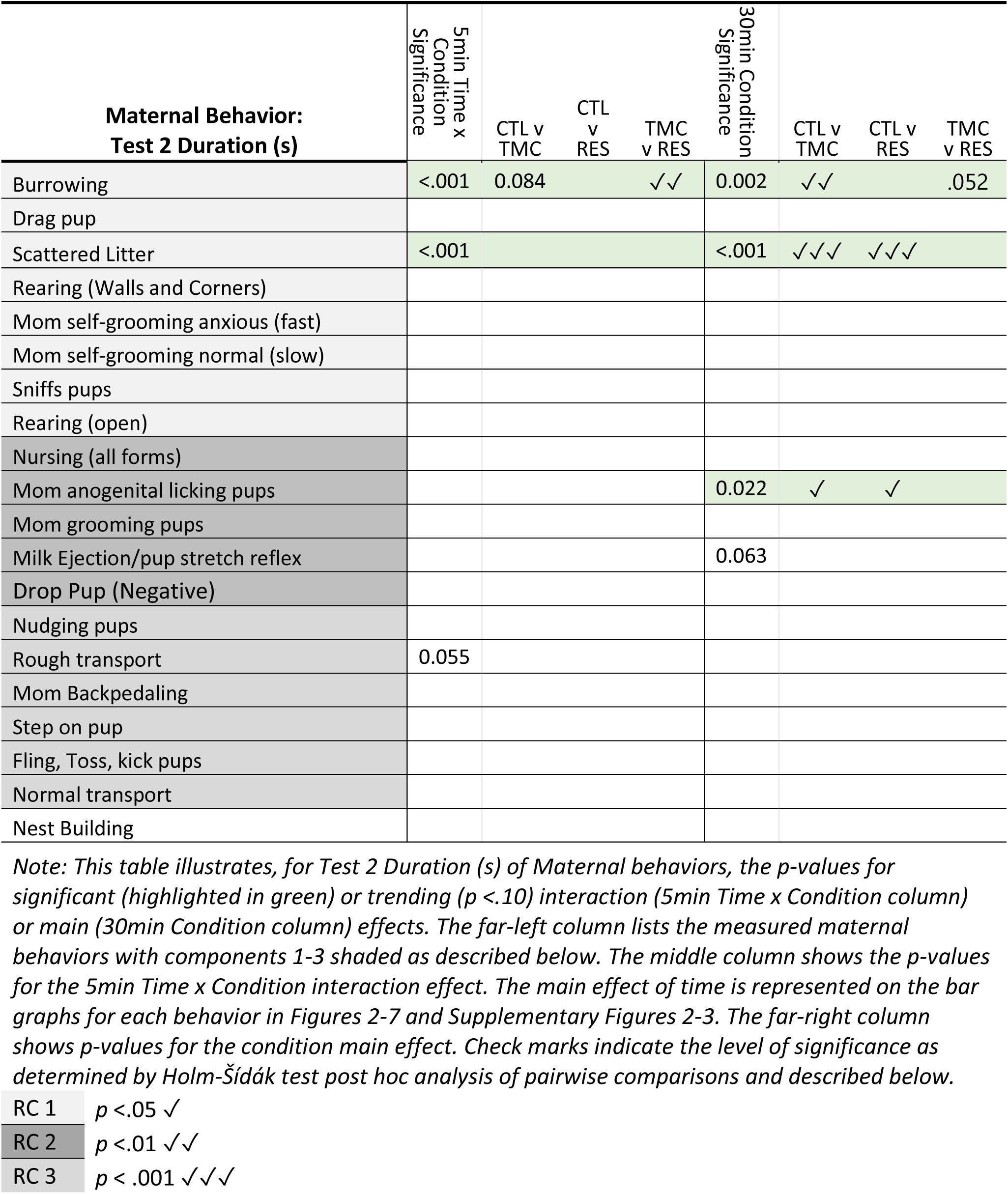
Summary of 2-way repeated measures ANOVA results from Day 5 (Test 2): Duration (s)

| Maternal Behavior:<br>Test 2 Duration (s) | 5min Time x<br>Condition<br>Significance | CTL v<br>TMC | CTL<br>v<br>RES | TMC<br>v<br>RES | 30min Condition<br>Significance | CTL v<br>TMC | CTL v<br>RES | TMC<br>v<br>RES |
| --- | --- | --- | --- | --- | --- | --- | --- | --- |
| Burrowing | <.001 | 0.084 |  | ✓✓ | 0.002 | ✓✓ |  | .052 |
| Drag pup |  |  |  |  |  |  |  |  |
| Scattered Litter | <.001 |  |  |  | <.001 | ✓✓✓ | ✓✓✓ |  |
| Rearing (Walls and Corners) |  |  |  |  |  |  |  |  |
| Mom self-grooming anxious (fast) |  |  |  |  |  |  |  |  |
| Mom self-grooming normal (slow) |  |  |  |  |  |  |  |  |
| Sniffs pups |  |  |  |  |  |  |  |  |
| Rearing (open) |  |  |  |  |  |  |  |  |
| Nursing (all forms) |  |  |  |  |  |  |  |  |
| Mom anogenital licking pups |  |  |  |  | 0.022 | ✓ | ✓ |  |
| Mom grooming pups |  |  |  |  |  |  |  |  |
| Milk Ejection/pup stretch reflex |  |  |  |  | 0.063 |  |  |  |
| Drop Pup (Negative) |  |  |  |  |  |  |  |  |
| Nudging pups |  |  |  |  |  |  |  |  |
| Rough transport | 0.055 |  |  |  |  |  |  |  |
| Mom Backpedaling |  |  |  |  |  |  |  |  |
| Step on pup |  |  |  |  |  |  |  |  |
| Fling, Toss, kick pups |  |  |  |  |  |  |  |  |
| Normal transport |  |  |  |  |  |  |  |  |
| Nest Building |  |  |  |  |  |  |  |  |

**Table 6.** Summary of 2-way repeated measures ANOVA results from Day 5 (Test 2): Frequency (s)

| **Maternal Behavior:**<br>**Test 2 Frequency (#)** | **5min Time x**<br>**Condition**<br>**Significance** | **CTL v**<br>**TMC** | **CTL v**<br>**RES** | **TMC v**<br>**RES** | **30min Condition**<br>**Significance** | **CTL**<br>**v**<br>**TMC** | **CTL v**<br>**RES** | **TMC v**<br>**RES** |
| --- | --- | --- | --- | --- | --- | --- | --- | --- |
| Burrowing | <.001 |  | ✓✓✓ | ✓✓✓ | 0.008 | ✓ |  | ✓ |
| Drag pup |  |  |  |  |  |  |  |  |
| Scattered Litter | 0.002 |  |  |  | 0.006 | ✓✓ | ✓✓ |  |
| Rearing (Walls and Corners) |  |  |  |  |  |  |  |  |
| Mom self-grooming anxious (fast) |  |  |  |  |  |  |  |  |
| Mom self-grooming normal (slow) |  |  |  |  |  |  |  |  |
| Sniffs pups | <.001 | ✓✓ | ✓✓ |  |  |  |  |  |
| Rearing (open) |  |  |  |  |  |  |  |  |
| Nursing (all forms) |  |  |  |  |  |  |  |  |
| Mom anogenital licking pups |  |  |  |  | 0.073 |  |  |  |
| Milk Ejection/pup stretch reflex | 0.012 |  |  |  | 0.097 |  |  |  |
| Mom grooming pups |  |  |  |  |  |  |  |  |
| Drop Pup (Negative) | 0.085 |  |  |  | 0.071 |  |  |  |
| Nudging pups | 0.048 |  | ✓✓ |  |  |  |  |  |
| Rough transport |  |  |  |  |  |  |  |  |
| Mom Backpedaling | 0.074 |  |  |  |  |  |  |  |
| Step on pup | 0.019 | 0.072 | ✓✓✓ | 0.072 |  |  |  |  |
| Fling, Toss, kick pups | 0.016 |  | ✓ | ✓✓ | 0.059 |  |  |  |
| Normal transport |  |  |  |  |  |  |  |  |
| Nest Building |  |  |  |  |  |  |  |  |

### RC1: Dam self-focused behaviors

Burrowing (Figures 2 and 5):

*First day.* Main effects of condition following the first day of testing indicate that during the entire 30min test, both the frequency *F*(2, 25) = 9.9, *p<*.001 and duration *F*(2, 25) = 7.0, *p=*.004 of burrowing was increased in the RES group relative to TMC and CTL. There was also a significant interaction of time and maternal condition for both duration *F*(10, 125) = 3.8, *p<*.001 and frequency *F*(10, 125) = 4.5, *p<*.001. During minute 0-5 and 5-10 of test, duration and frequency of manipulating cage bedding was significantly increased in RES relative to TMC and CTL.

*Fifth day. F*ollowing the fifth day, there was main effect of condition for both duration [*F*(2, 25) = 8.4, *p=*.002] and frequency [*F*(2, 25) = 5.9, *p=*.008] of burrowing; over the entire 30min, CTL and RES had higher duration and frequency of burrowing relative to TMC. There was a significant time and condition interaction for duration [*F*(10, 125) = 3.6, *p<*.001] and frequency *F*(10, 125) = 5.9, *p<*.001. During minutes 0-5, duration of burrowing was higher in RES than TMC and CTL. During min 5-10 both RES and TMC were higher than TMC. During min 15-20 CTL was higher than TMC and RES. For Frequency, during min 0-5, RES was higher than TMC and CTL. During min 5-10, both RES and CTL were higher than TMC and during min 15-20 CTL was higher than TMC and RES.

Dragging nipple-attached pup (Supplemental Figures 2 and 3):

*First and Fifth days.* There was no significant difference in the frequency or duration of dragging nipple-attached pup between RES, TMC and CTL dams (*p’*s > .05) at any time during the testing period.

Scattered Litter (Figures 2 and 5):

*First day.* There was no significant difference in scattered litter frequency or duration between RES, TMC, and CTL dams *p*> .05) at any time during the test.

*Fifth day.* In contrast, following the fifth day, the main effect of condition revealed that the frequency [*F*(2, 23) = 6.3, *p=*.006] of the litter being scattered decreased in CTL relative to TMC and RES but the duration [*F*(2, 24) = 13.0, *p<*.001] of litter being scattered was increased in CTL relative to RES and TMC. There was an interaction of time and condition for frequency [*F*(10, 115) = 3.0, *p=*.002] and duration [*F*(10, 120) = 5.7, *p<*.001]. During min 10-15, 15-20, 20-25, and 25-30 duration of CTL scattered litter was higher than RES or TMC. During min 20-25 and 25-30, frequency of scattered litter was lower in CTL relative to RES and TMC.

Rearing (Walls and Corners, Figures 2 and 5):

*First and Fifth days.* No significant differences, on the first or fifth day, in wall/corner rearing frequency or duration between RES, TMC and CTL dams *p*’s > .05) at any time during the test.

Mom self-grooming anxious (fast, Figures 2 and 5):

*First day.* Main effect of condition suggest that during the 30 min test, both the frequency *F*(2, 24) = 4.1, *p=*.029 and duration *F*(2, 24) = 3.5, *p=*.046 of dam self-grooming fast was increased in RES relative to TMC and CTL although post-hoc analyses for frequency and duration failed to reach significance (*p’*s > .057).

*Fifth day.* In contrast, following the fifth day, there were no observed instances of self-grooming (fast) during test 2.

Mom self-grooming normal (slow, Supplemental Figures 2 and 3):

First and Fifth days. There were no significant differences in mom self-grooming slow frequency or duration between RES, TMC and CTL dams *p*’s > .05) at any time during the tests.

Sniffs pups (Figures 2 and 5):

*First day.* There was no significant difference in dams sniffing pups frequency or duration between RES, TMC and CTL dams *p*’s > .05) at any time during the test

*Fifth day.* There were no main effect of condition but there was a significant interaction of time and condition on sniffing frequency [*F*(10, 130) = 3.3, *p*<.001]; during minutes 0-5, sniffing was higher lower in CTL than RES and TMC.

### Rearing (Open [Supplemental Figures 2 and 3])

*First and Fifth days.* No significant differences, on the first or fifth day, in rearing in the open frequency or duration between RES, TMC and CTL dams *p’*s > .05) were observed at any time during the test.

### RC2: Nurturing/typical maternal behavior

Nursing (Figures 3 and 6):

*First day.* For the entire 30min test the duration *F*(2, 26) = 5.4, *p=*.011 (but not frequency, p > .05) of nursing was significantly decreased in RES relative to CTL (but not TMC).

*Fifth day.* There were no significant differences in nursing frequency or duration between RES, TMC, and CTL at any time point (all *p’*s > .05).

Dam Anogenital licking pups (Figures 3 and 6):

*First day.* Following the first day, dams showed no significant difference in pup anogenital licking frequency or duration between RES, TMC and CTL groups (*p’*s > .05) at any time during the test. *Fifth day.* a main effect of condition revealed decreased duration [*F*(2, 26) = 4.4, *p=*.022] but not frequency [*F*(2, 26) = 2.9, *p=*.073] of anogenital licking on pups in CTL relative to RES and TMC during the entire 30min test. No other significant effects were found.

Dam grooming pups (Supplemental Figures 2 and 3):

*First and Fifth days.* There was no significant difference in grooming pups frequency or duration between RES, TMC and CTL dams *p’*s > .05) at either time during the test.

Milk Ejection/pup stretch reflex (Figures 3 and 6):

*First day.* There was no significant difference in milk ejection frequency or duration between RES, TMC, and CTL dams (*p’*s > .05) at any time during the test (suggesting that even with overall shorter duration of nursing, pups still received similar nourishment across conditions).

*Fifth day.* There was no main effect of condition but there was an interaction between time and condition for milk ejection frequency [*F*(10, 130) = 2.4, *p=*.012] but not duration [*F*(10, 125) = 1.3, *p=*.265]. During minutes 20-25 and 25-30 milk ejections were higher in RES than TMC and CTL. During min 25-30, RES was higher than CTL and TMC, but TMC was also significantly higher than CTL.

Drop Pup during transport (Negative loading, Supplemental Figures 2 and 3):

*First and Fifth days.* No significant differences, on the first or fifth day, in drop pup frequency between RES, TMC, and CTL dams *p>* .05) were observed at any time during the test.

### RC3: Rough Treatment of Pups

Nudging pups (Figures 4 and 7):

*First day.* There was no significant difference in dams nudging pups frequency or duration between RES, TMC, and CTL dams *p’*s > .05) at any time during the test.

*Fifth day.* Similarly, there was no main effect of condition, although there was a time x condition effect on frequency [*F*(10, 130) = 1.9, *p=*.048] such that during minutes 0-5, RES nudging was higher than CTL, and during minutes 5-10, RES and TMC nudging were higher than CTL.

Rough pup transport (Figures 4 and 7):

*First day.* Main effects of condition and post hoc indicate that during the entire 30min test, pup rough handling was significantly increased in the RES group relative to TMC and CTL in frequency *F*(2, 26) = 8.4, *p=*.002, and RES relative to CTL in duration *F*(2, 25) = 7.7, *p=*.002. Incidence of rough transport of the pups showed a significant interaction of time and maternal condition for both duration *F*(10, 125) = 3.5, *p*<.001 and frequency *F*(10, 130) = 2.3, *p=*.018. During minutes 0-5 of test, duration of rough transport was longest in RES, significantly reduced in TMC, and lowest in CTL dams. Similarly, frequency of rough transport during minutes 0-5 was significantly higher for RES, with TMC and CTL not differing. During minutes 5-10, RES rough transport frequency remained elevated above CTL and trending (*p* =.07) above TMC.

*Fifth day.* Differences in rough transport frequency or duration between RES, TMC and CTL dams (*p’*s > .05) were not observed at any time during the test following the fifth day.

Backpedaling during pup transport (Supplemental Figures 2 and 3):

*First and Fifth days.* No significant differences, on the first or fifth day, in dam backpedaling frequency or duration between RES, TMC and CTL dams *p’*s > .05) were observed at any time during the test.

Step on pup (Negative loading RC2, Figures 4 and 7):

*First day.* There was no significant difference in the duration of time stepping on pups between conditions (p >.05). However, over the entire 30min test, the frequency *F*(2, 27) = 4.7, *p=*.018 of stepping on pups was significantly lower in the CTL group relative to both RES and TMC groups. Further, an interaction between time and condition revealed that this increase was most pronounced during minutes 0-5 when the frequency of stepping on pups was highest in TMC, lower in RES, and lowest in CTL.

*Fifth day.* Following the fifth day, there was no main effect of condition on stepping on pups but there was a time x condition effect of frequency [*F*(10, 130) = 2.2, *p=*.019] such that stepping on pups was higher in RES relative to CTL during minutes 0-5.

Fling, Toss, kick pups (Figures 4 and 7):

*First day.* Main effects of condition indicate that during the entire 30min test, both the frequency *F*(2, 24) = 11.0, *p*<.001 and duration *F*(2, 24) = 13.0, *p*<.001 of flinging/tossing/kicking pups was significantly increased in the RES group relative to TMC and CTL. There was also a significant interaction of time and maternal condition for both duration *F*(10, 120) = 6.7, *p*<.001 and frequency *F*(10, 120) = 7.5, *p*<.001. During minutes 0-5 and 5-10 of test, duration and frequency of flinging/tossing/kicking was higher in RES relative to TMC and CTL.

*Fifth day.* Similarly, although there was no main effect of condition, there was a time x condition effect on frequency [*F*(10, 130) = 2.3, *p=*.016] such that during minutes 0-5 and 5-10, RES flinging was higher than CTL and TMC.

Normal pup transport (Figures 4 and 7):

*First day.* Main effects of condition indicate that during the entire 30min test, both the frequency *F*(2, 26) = 4.6, *p=*.019 and duration *F*(2, 27) = 3.7, *p=*.038 of normal transport was reduced in the RES group relative to TMC and CTL. However, for duration, post-hoc test revealed this decrease was not significant between RES and TMC (p = 0.069) and RES and CTL (p = 0.078). There was also a significant interaction of time and maternal condition for both duration *F*(10, 135) = 3.1, *p=*.002 and frequency *F*(10, 130) = 3.1, *p=*.002. Post-hoc revealed during minute 0-5 of test, duration and frequency of normal transport was significantly reduced in RES relative to TMC and CTL.

*Fifth day.* In contrast, following the fifth day, there were no significant differences in normal transport frequency or duration between RES, TMC, and CTL at any time point (all *p’*s > .05).

### Behavior not loading on the three components

Nest Building (Supplemental Figures 2 and 3):

*First and Fifth days.* No significant differences, on the first or fifth day, in nest building frequency or duration between RES, TMC and CTL dams (*p’*s > .05) were observed at any time during the test.

### Maternal changes in regional cFos activity

Fos expression following Day 5 (Figures 8-9)

<u>mPFC</u> (Figure 8). In the infralimbic mPFC, there was a significant effect *F*(2,21)=11.50; *p*<0.001). Post hoc tests indicated that both the TMC (*p*<0.001) and the RES (*p*<0.004) groups exhibited higher number of cFos-expressing cells than the CTL mothers. In the prelimbic mPFC, there was a significant effect *F*(2,21)=9.48; *p*<0.004) in the PL. Post hoc tests indicated that both the TMC (*p*<0.002) and the RES (*p*<0.01) groups exhibited higher number of cFos-expressing cells than the CTL mothers.

<u>BLA</u> (Figure 8). For the BLA, there was a significant effect *F(*2,16)=3.89; *p*<0.042). Post hoc tests indicated that the number of cFos-expressing cells in the BLA was significantly higher in the RES mothers compared to the CTL mothers (*p*<0.03). There was no difference between the TMC and RES moms or between TMC and CTL mothers.

<u>MEA</u> (Figure 8). For the MEA, there was a significant effect *F(*2,15)=8.21; *p*<0.001). Post hoc tests indicated that the number of cFos-expressing cells in the MEA was significantly higher in the RES mums compared to the CTL mothers (*p*<0.001) and tended to be significant compared to the TMC mothers (*p*<0.06).

<u>PAG</u> (Figure 9). In the dorsomedial PAG (dm PAG), there was a significant effect *F(*2,18)=5.24; *p<*0.016). Post hoc tests indicated that the number of cFos-expressing cells in the dmPAG was significantly higher in the RES mothers compared to both the CTL (*p<*0.01) and TMC (*p<*0.03) mothers. In the dorsolateral PAG (dlPAG), there was a significant effect *F(*2,19)=14.3; *p<*0.001). Post hoc tests indicated that the number of cFos-expressing cells in the dlPAG was significantly higher in the RES mums compared to both the CTL (*p<*0.001) and TMC (*p<*0.001) mothers. In the Lateral PAG, there was no significant differences in the number of cFos-expressing cells between groups.

<u>PVT</u> (Figure 9). For the PVT, there was a trend towards significance (*p<*0.08). Perusal of the graphs suggests that naïve and control groups trended to be different from one another.

### Stress-related corticosterone responses in post-weaning Dams

There was a significant Time effect *F(*3,132)=72.7; *p<*0.001) and a significant Group effect (F2,132)=13.41; *p<*0.001) but no significant interaction effects. Post hoc results comparing groups at a given time point are most relevant. At 0min (baseline), there was no difference between groups. At 15min, naïve groups exhibited higher corticosterone compared to the RES mothers (*p<*0.02). At 30min, RES groups were higher than both CTL and TMC mothers (<0.001 and 0.0099, respectively) and CTL and TMC were not different from one another. At 60min, RES mothers were lower than TMC (*p<*0.05). Integrated corticosterone analysis showed that there was a significant main effect *F(*2,34)=4.99; *p<*0.01) with post-hocs indicating that RES mothers exhibited lower corticosterone compared to CTL mothers (*p<*0.01).

### Forced swim test

No significant differences were observed in time spent immobile, swimming or climbing between groups (all *p’*s >.05). No significant differences were observed in latency to become immobile (all *p’*s >.05).

## Discussion

Our data identified both immediate and sustained alterations in maternal behavior and activation of specified brain regions associated with acute or repeated postpartum social stress for the dam. Following a single exposure to an intruder, nurturing maternal behaviors were reduced, maternal rough behaviors significantly increased, and maternal stress (e.g., burrowing and self-grooming) behaviors elevated. Importantly, after five days of social stress, some behavioral changes return to control levels, but distinct patterns of change in behaviors and neural activity remain, suggesting a unique stress effect of repeatedly defeating an intruder compared to repeated separation from their pups or separation alone. There are also consequences for maternal behavior when pups are removed from the dam without the intruder stress, suggesting multiple but distinct effects of different types or intensities of stressors on maternal stress neurobiology.

### The social stress of fighting an intruder alters maternal behavior

The balance between approach and withdrawal behaviors towards pups naturally shifts during different phases of maternal care. Initially, approach behaviors dominate immediately after birth and are maintained through the middle of the preweaning period before ending with predominance of withdrawal/avoidance up to weaning^19^. In addition, maternal aggression is a unique form of aggression that occurs under the requisite external environment (including the pups to be defended) and an internal neuroendocrine environment that typically follows birth ^20–22^. Successful rearing necessitates that the hormonal and neural bases of this behavioral cascade produce appropriately nurturing maternal care.

Acute threats during the postpartum period shift behavioral choice from pup-directed maternal care to pup-protecting aggressive and defensive behaviors. When dams and their pups are threatened, maternal behaviors like nursing and pup grooming are reduced. Then, aggression towards an intruder subsequently induces more defensive burrowing and self-grooming. Although this is particularly pronounced when the pups remain in the home cage during the threat exposure ^43,45^, here we show that maternal aggression/defense persists even in the absence of an imminent threat to the pups. Unsurprisingly, following the first intruder exposure, nursing decreased and stress-related behavior (e.g. anxious self-grooming and burrowing) trended high in RES dams relative to dams that were separated at the same time without defeat stress or not separated at all. Whereas anxious self-grooming diminished by Day 5, the occurrence of abusive handling (including roughly transport and kicking pups) and defensive burrowing in RES dams remained high between from Day 1 (PD7) and Day 5 (PD11). This pattern of generalized defensive behaviors to acute environmental stressors (like predator odor and electrified rods ^45,62,63^) has been previously described. For rats, when an electrified shock grid is encountered in the home cage, dams retrieve and relocate the pups to new nest areas and attempt to bury the rod by burrowing through and moving bedding toward the probe ^62,63^. This previously described active coping strategy is mediated by the central amygdala ^64–66^. Our data suggest that, after the intruder is removed following a single-day acute exposure, targeted aggression/defensive maternal behaviors become more generalized, with increases in active coping and anxiety behaviors that may make the dam less attentive to pups when handling them by redirecting her attention more generally to the environment.

Following five days of social stress, some behaviors acutely disrupted in RES dams return to control levels, suggesting altered strategies for maternal behavioral choice in response to the threat. Generally, maternal care and aggressive behavior peak after the first postnatal week, and naturally wane between the second and third postnatal weeks ^19,20,67,68^. However, Nephew and colleagues ^13,15,57^ suggest that repeated postpartum social stress may sensitize maternal aggression, causing its prolonged expression at the expense of maternal care behaviors. They demonstrated that while maternal aggression in control dams decreased between PD9 and PD16, aggression remained elevated in RES rats exposed to repeated social stress from PD2-16 ^13,57^. Other maternal behaviors were also altered in their studies between PD2 and PD9; stress increased pup retrieval, nesting (sometimes relocating the nest entirely), self-grooming, and activity/patrolling while decreasing nursing and pup grooming ^13,15^ suggesting that dams forgo pup-directed maternal care in exchange for threat-directed or self-directed care. However, they also show that maternal anxiety-like behaviors and most maternal care behaviors, like nursing, pup grooming, and nesting, returned to control levels by PD16. Similarly, our results show that dam self-grooming, the total duration of nursing, and pup transport returned to control levels by PD11 (Day 5 of testing), but defensive burrowing and flinging/kicking pups remained elevated in RES dams, demonstrating a similar normalization of maternal care but not maternal aggressive/defensive behaviors following repeated social stress.

Physiological changes (e.g. HPA response to stress and nursing behavior) in dams were evident across multiple measures in the current report and similar findings have been reported in other studies. Reduced maternal nursing efficiency may be a consequence of postpartum social stress. Although RES dams increased nursing relative to controls, their pups showed similar growth between the groups, suggesting a physiological inefficiency in their nursing ^15^. We similarly found that at Day 5 testing, there was an increase in frequency of RES dam milk ejections but no difference in total nursing duration, suggesting a compensatory increase in some aspects of nursing that may support ongoing growth of the pups. By Day 5 of testing, maternal anxious self-grooming durations were similar between RES and the control groups. However, assessment of CORT levels after weaning (10 days after the last intruder exposure) revealed reduced CORT concentrations following restraint stress in RES relative to controls, indicative of a disruption of HPA function. It is possible that the decrease in stress behavior observed on Day 5 in our report is due to compensatory down-regulation of stress systems through a mechanism typically activated to regulate maternal response to stressors during lactation ^69^. Characteristically, basal CORT levels are elevated during lactation; however stress-induced CORT expression is blunted relative to controls ^25,70,71^. This downshift in CORT may be designed to facilitate the rapid return to maternal care following a stressful event but is aberrantly maintained following repeated stress exposures. Additional testing is needed to explore this possibility and how it interacts with midbrain and corticolimbic control of maternal behavior expression.

### Following pup separation, the medial prefrontal cortex may drive pup-directed attention

Following the end of the maternal manipulations on Day 5, both the RES and TMC dams showed increased mPFC activation as assessed by Fos expression (in both the prelimbic and infralimbic subregions). This suggests that the mPFC is most strongly responding to pup separation, rather than to social stress induced by intruder threat. The mPFC may support several aspects of maternal behavior, although it is most clear that mPFC is likely playing an executive control function to organize maternal behavior; mPFC disruption typically impairs or delays maternal behavior, but, importantly, does not completely eliminate it ^31–37^.

There are different roles of the IL and PL in response to exposure to stress and response to pups. Inactivation of dam IL on postpartum day 8 biased conditioned place preference (CPP) away from the cocaine-associated compartment and toward the pup-associated compartment whereas inactivation of the PL had the opposite effect^35^. The authors suggest that, rather than playing a direct role in processing the motivational value of pups, IL resolves conflict to maintain maternal behavior during times of competing motivations. This suggests that PL does the opposite, shifts focus away from pups and toward the competing motivation, perhaps to activate aggression/protective behaviors. But when there was no conflict, inactivation had no effect on preference in IL or PL groups. However, this lack of PL and IL effect on maternal behavior following inactivation contrasts with other studies. Asina-Llanes and Olazábal^72^ suggest that the role of the mPFC (particularly the PL of the mPFC) is the rapid induction of maternal behavior by reducing anxiety toward pup-related cues. Excitotoxic lesions in their females delayed the onset of full maternal behaviors in pup-naive virgin mice but did not change infanticidal behavior (which they speculate is a more impulsive, non-mPFC-mediated behavior). Also, lesioned animals did not retrieve pups in the first 15min (100% of control dams retrieved in the first 15min); however, licking bouts were not different between groups, demonstrating dissociable behavior patterns.

In our data, concurrent activation of IL and PL in both RES and TMC likely does not reflect conflict resolution; the dams are alone with pups following testing for both groups. Rather, increased activation in both subregions may reflect natural transitional control of maternal behavior from IL- to PL-dominant. Wu et al.,^73^ suggest that the mPFC facilitates the transition to maternal behavior through its projections to the mPOA during lactation and increases pup-directed maternal behaviors (e.g. pup retrieval from outside to inside the nest) in the presence of a threat. Although Wu et al., ^73^ did not distinguish between IL and PL neuronal populations, maternal nurturing behavior is driven by IL in early lactation and maintained by PL in late parturition (for review, see ^74^). PD11 (Day 5 of testing in our procedure) is a transitional period for pup activity ^75^ and it may be a transitional period in dams, where behavioral choice is a shared activity between IL and PL as dams shift from nursing the mostly nest-bound mound of young pups to retrieving and protecting older pups that are beginning to explore outside the nest. If this shift is indicative of a transitional neurobehavioral state, we may expect that repeated social stress during an earlier window like PD4-9 may show stronger effects on maternal IL activity relative to dams at PD10-15 or later windows that will likely affect PL activity. Alternatively, different subpopulations of IL and PL neuron may control dissociable aspects of the maternal behavior. Further investigation is required to dissociate the distinct contribution of PL and IL to the behavioral changes observed in the current report.

### Amygdala activity but may reflect modulated emotional behavior expression between groups

Fos expression in the medial and basolateral amygdala differed from that of the PFC or PAG in that there were differences between the RES and other groups, depending on the amygdaloid subregion. The BLA is classically considered the “input” site and receives information from all sensory systems that is preprocessed by the insula or cingulate cortices and the sensory thalamus, VTA and PAG ^76–80^. The BLA then activates the neuronal activity that is required for fear acquisition and appropriate responses ^124^. Likewise, the MeA receives input from multiple sites including the olfactory system through pheromones, as well as from crucial areas such as the hypothalamus, hippocampus, and reward system. In our paradigm, activation of these amygdala nuclei was selective to intruder stress relative to CTL and appeared reduced but not statistically different (*p=*.06) from TMC. That the distinction in activity between RES and TMC was not stronger, specifically in the BLA, was surprising. While both nuclei of the amygdala receive olfactory input to help process pup odors and have reciprocal connections with the mPOA^16,81–83^, the BLA is more linked to maternal aggression towards a threat to her pups.

Whether this response pattern of the amygdala indicates mixed responding to the different behavioral demands of pup separation relative to pup protection or modulated negative emotional states between the three groups (or some other process) requires additional investigation. An interesting possibility is that the amygdala is more grossly activated by intruder exposure in RES, but the overall stress of pup removal in TMC still triggers some Amygdala activation; these two experiences are then disambiguated by the PAG (see below).

This would demonstrate an AMG-PAG relationship whereby coarse stress-related modulation of maternal behavior is fine-tuned by the PAG and the distinct behavioral patterns observed in our RES groups require both systems to be repeatedly activated.

### The PAG is reactive strictly to the aggressive encounter

The dorsal and dorsolateral PAG serves as an integration hub for both maternal care and maternal aggression/threat responding and relays input from rostral regions to caudal sites, including spinal cord ^16,84^. There is a demonstrated association between PAG function and maternal behaviors in rodents wherein PAG contributes to behavioral selection (pup retrieval, nest building, nursing) in lactating dams, integrating hormonal status and input from the mPOA and downstream sites differentially along its dorso-ventral axis. *Activation* of rostral-lateral (rl)PAG opioid receptors can impair maternal behaviors like pup retrieval by *increasing latency* to retrieve pups when dams have to choose between pup retrieval and predation of cockroaches ^49^. The opposite effect is observed following rlPAG inhibition; disrupting rlPAG via *lesioning* significantly *reduced latency* to first pup retrieval in dams with pups exposed to cat odor relative to intact dams exposed to cat odor ^45^. The dorsal (d)PAG has a contrasting profile of behavioral change to that of the rlPAG. Serotonin reduces PAG activity, with the strongest inhibition observed in dorsal PAG regions ^85,86^. However, in contrast to rlPAG disruption, dPAG inhibition through increased serotonergic activity reduces maternal aggression toward intruders in 7-day postpartum rats but pup care, including retrieval, was unaffected ^46–48^. In the current report, dmPAG and dlPAG showed increased activation in RES dams relative to TMC and CTL after 5 days of defeating an intruder, reflecting recruitment of intruder-specific maternal defense circuitry that is not activated simply by removing the pups (i.e., the TMC dams). lPAG did not show this difference, suggesting that scattering pups (which happened in all three groups) similarly activated maternal retrieval circuits.

Importantly, the contrast between dorsal PAG and lateral PAG activation emphasizes the distinct roles of the PAG subregions and suggests that dysregulated dPAG activity, following repeated social stress, may support the sensitized/sustained maternal aggression state previously suggested ^13,14^ and demonstrated here. Our data suggests that this shift in maternal behavior and long-term effects on maternal stress responding and (at least in part) mediated by the dPAG. Further, it is likely that targeting this circuit and its connections to the amygdala or mPFC subregions during earlier (before PD10) or later lactation periods may reveal additional distinct roles of the amygdala and mPFC subregions and whether they differentially (or at all) contribute to the shifts in behavioral expression observed in dams following repeated social stress.

To our knowledge, ours is the first PCA to statistically characterize this maternal “anxious-inattentive” behavioral profile, with RC1 (containing dam defensive burrowing, anxious grooming) and RC3 (containing flinging/kicking, stepping on, and rough transport pups) showing significant RES effects relative to controls. Distinct from previous work, our PCA also indicates that behaviors within a component tend to change in similar ways between Day 1 and Day 5 of testing. This may reflect a common neurobiological substrate responsible for shifts in specific component behaviors. Indeed, dysregulation of AMG-PAG relationship (perhaps irrespective of mPFC contributions) likely facilitates the distinct pattern of RES behavioral changes wherein repeated defeat stress sensitizes AMG activity (Figure 8) that is further accentuated by PAG increases, thereby sustaining defensive-related anxiety and reducing nurturing maternal behavior. Manipulation of this microcircuit will reveal whether sustained alteration of maternal behavior following repeated social stress is controlled by dysregulation in this PFC-AMG-PAG network.

### Future directions

We made an initial effort to define a limited number of brain sites involved in maternal behavior and maternal stress/aggression but clearly there are multiple brain regions and subregions that also are likely involved which we did not study. These might include other amygdala and hypothalamic nuclei, and limbic structures such as the nucleus accumbens. We also focused on test intervals immediately after the last day of intruder stress and do not know whether changes in behavior and neural circuit activation are long lasting and if so, do they influence subsequent episodes of pregnancy and maternal care. Likewise, we did not determine if the intruder stress altered subsequent responses to different stressors. For example, would the propensity to be vulnerable or resilient to later non-maternal stress be different amongst the different maternal groups. These questions on how maternal aggression subsequently impacts her behavior and that of her offspring remain unanswered. While addressing these possibilities was beyond the scope of the current manuscript, future experiments that systematically assess these other potential neurobehavioral components of maternal stress-related behavioral changes are an exciting future direction.

### Summary

We found that stressing the dam by intruder threat, alters subsequent behavior towards the pups and that the manner of stress matters. Some behaviors were affected simply by an hour separation from the pups (burrowing; nursing) but a larger effect was seen when the dam was threatened by an intruder after separation. In this latter case there was atypical, rougher treatment of the pups (flinging; rough transport) over and above that seen with simple separation. Following five days of stress, the effects of the intruder stress were partially mitigated in their duration and more typical maternal care began to appear. Our results indicated that the mPFC was activated primarily by pup separation whereas the amygdala and PAG were differentially activated by intruder threat, regardless of pup separation. These results suggest that different neural circuits underlie the maternal response to intruder threat than those that underlie the maternal response to pup separation. Studies to examine these circuits are underway.

## Funding Sources

Research reported in this publication was supported by the Eunice Kennedy Shriver National Institute of Child Health and Human Development of the National Institutes of Health under award #1L40HD120184-01 to PAR.

## Notes

### Competing Interest Statement

The authors have declared no competing interest.

