## Supplemental Figures for "Maternal defense against intruders changes her subsequent maternal behavior and neural circuitry"

_Supplemental Figure1. The Scree plot from the PCA analysis that shows that three components explained more variance than the simulated data and thus we focused on those first three components._

_Supplemental Figure-2-3: Additional maternal behaviors. Mean (±SEM) count of c-Fos expression cells in select regions of the brain. The group sizes for the CTL, TMC, and RES dam groups for each brain region is as follows: PVT n=11, 7, and 7; for PL and IL mPFC n=11, 7, and 6; MEA n=6, 5, and 7; for all PAG subregions n=11, 6, and 5. *indicates $p < .05$._

_Supplemental Figure-4-5: Estimation Plots for significant changes in maternal behaviors. These figures supplement each significant two-group comparison from Tables 3 and 4. Frequency data are shown as examples. Similar plots for the duration data are available on request. Each figure presents the data points (left panel) and mean difference between the two conditions plus/minus the standard deviation (right panel) for a single behavior. Day 1/Day 5 refer to the first or fifth defeat trial; 5 min/30 min refer to the first five minutes or the total 30-minute session of the dam's behavior upon return of her pups. The estimation plots largely confirm the ANOVA multiple comparison results._

**Supplemental Figure 1**: *Scree Plot of the PCA Analysis*

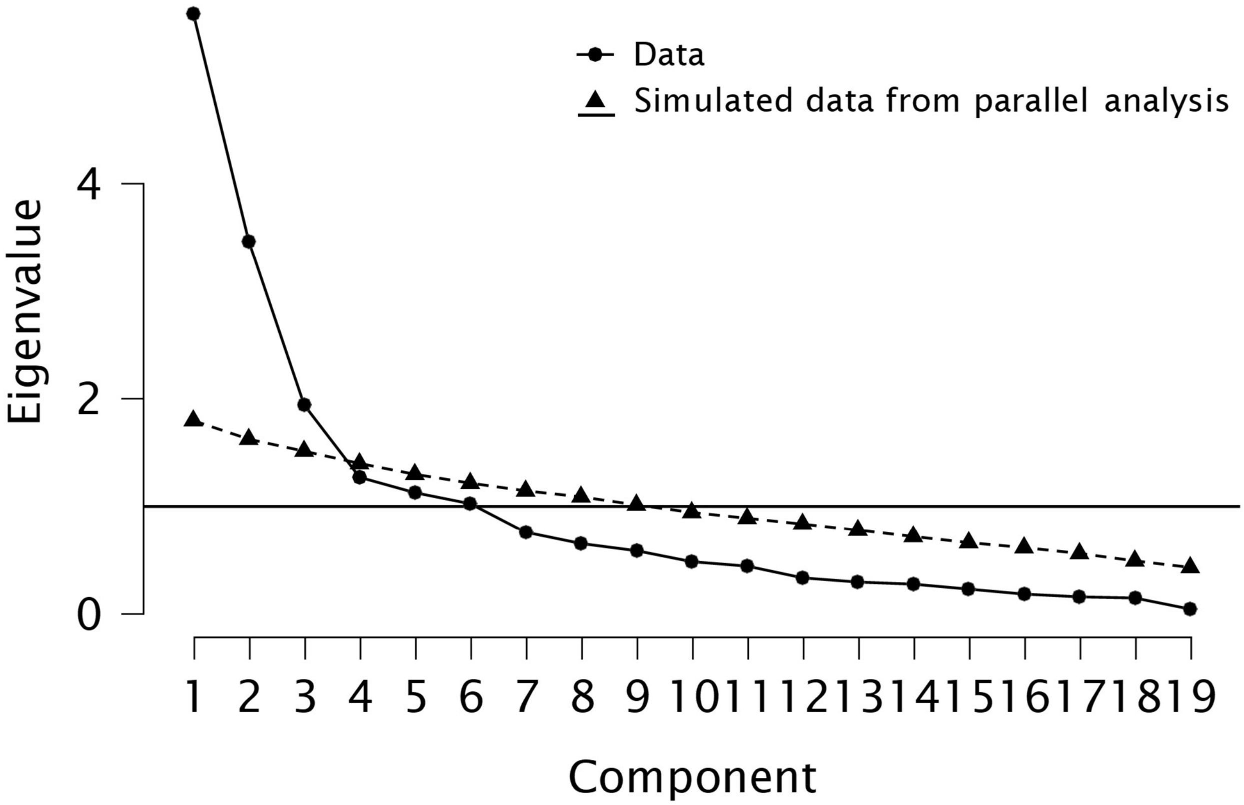

**Supplemental Figure 2**: *Maternal Behavior Day 1 (Test 1) Additional Behaviors*

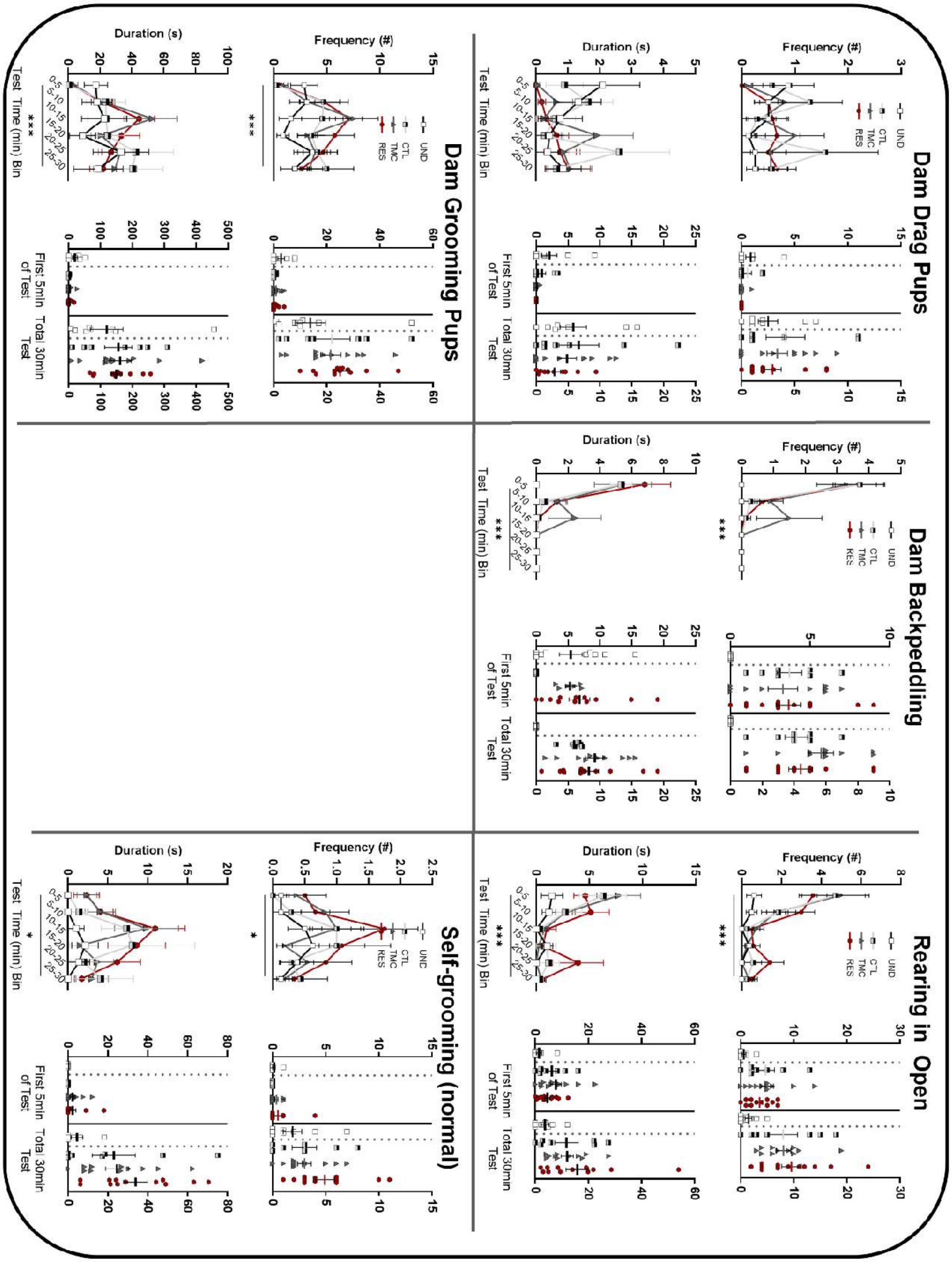

**Supplemental Figure 3:** *Maternal Behavior Day 5 (Test 2) Additional Behaviors*

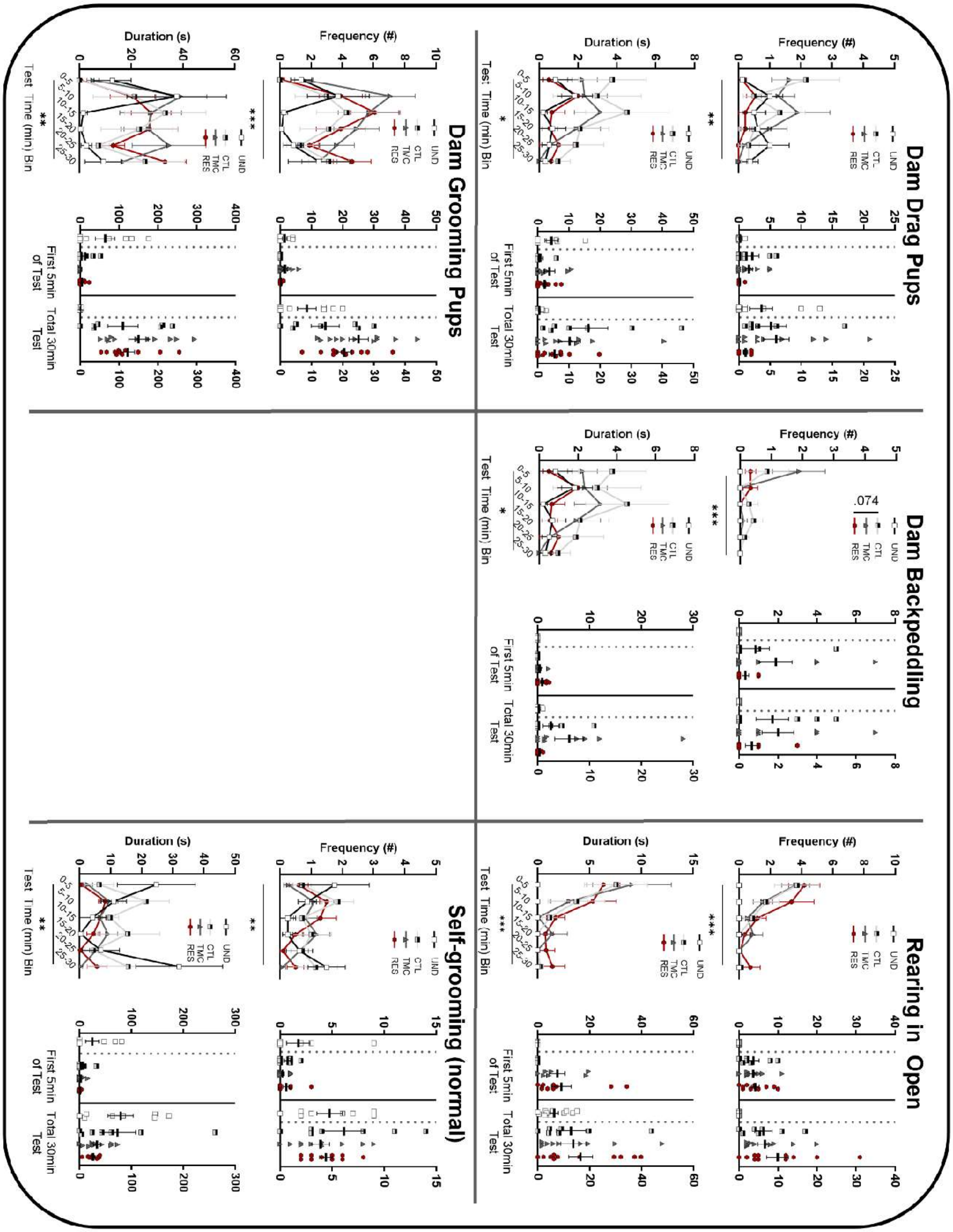

**Supplemental Figure 4:** *Estimation plots for significant changes in maternal behaviors*

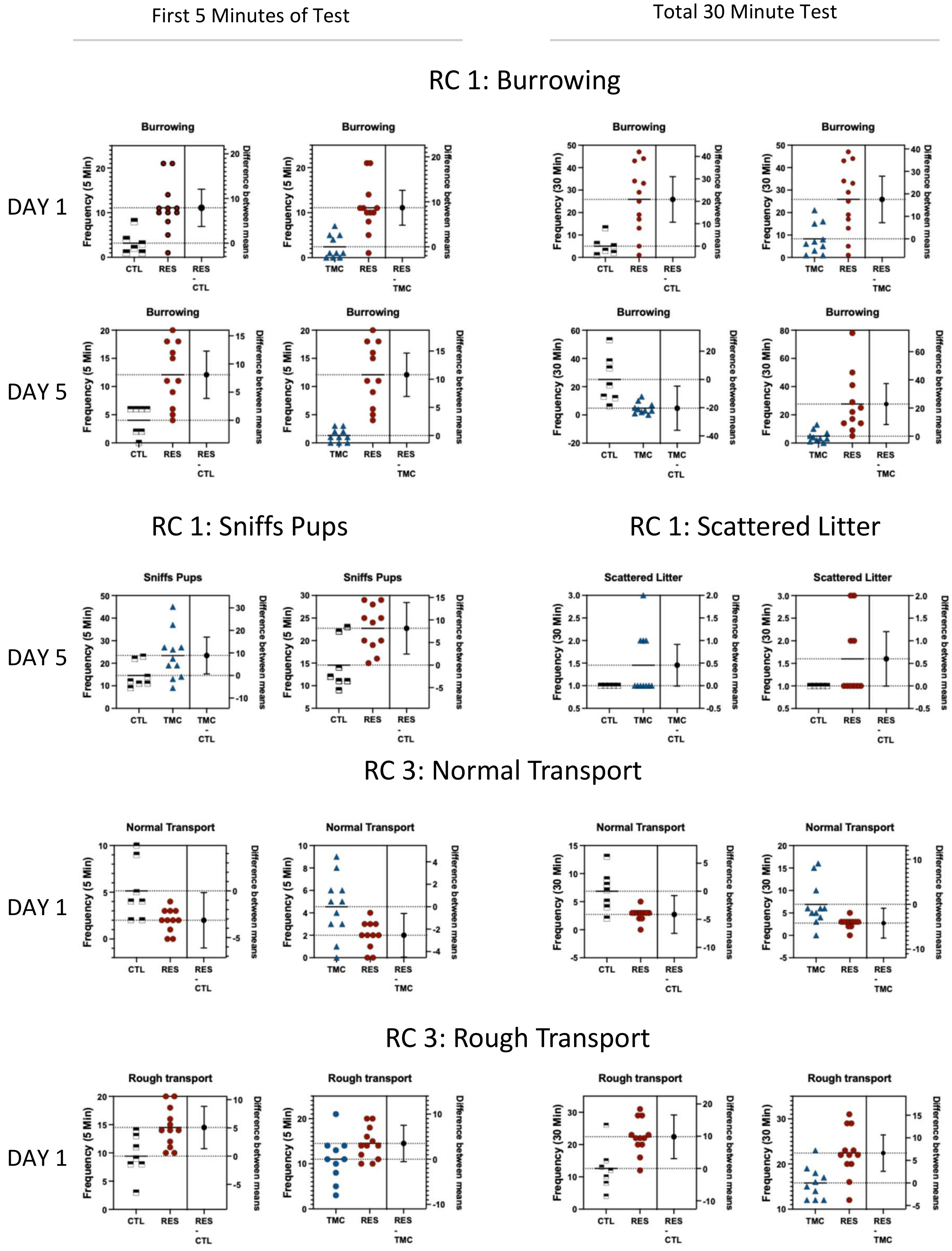

26 **Supplemental Figure 5:** *Estimation plots for significant changes in maternal behaviors (Cont'd)*

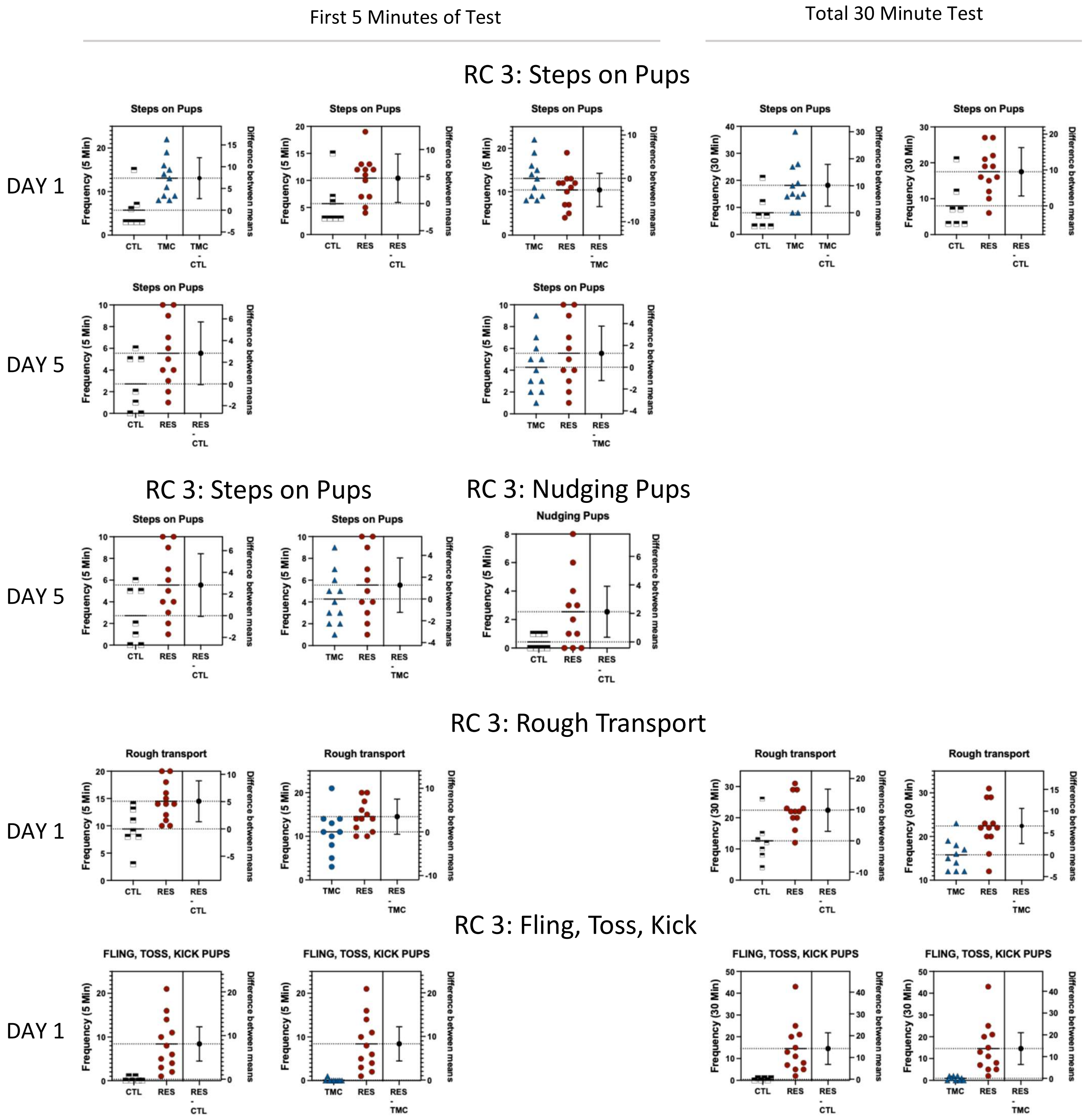
